# Capturing global pet dog gut microbial diversity and hundreds of near-finished bacterial genomes by using long-read metagenomics in a Shanghai cohort

**DOI:** 10.1101/2025.09.17.676595

**Authors:** Anna Cuscó, Yiqian Duan, Fernando Gil, Alexei Chklovski, Nithya Kruthi, Shaojun Pan, Sofia Forslund, Susanne Lau, Ulrike Löber, Xing-Ming Zhao, Luis Pedro Coelho

## Abstract

Pet dogs are considered part of the family, and understanding their gut microbiomes can provide insights into both animal and household health. Most comprehensive studies, however, relied on short-read sequencing, resulting in fragmented MAGs that miss mobile elements, antimicrobial-resistance genes, and ribosomal genes. Here, we applied deep long-read metagenomics (polished with short-reads) to fecal samples from 51 urban pet dogs in Shanghai, generating 2,676 MAGs—representing 320 bacterial species—, of which ∼72% achieved near-finished quality, often improving on the corresponding reference public genome. Comparisons with external datasets showed that our Shanghai-based MAG catalog is representative of pet dogs worldwide (median read mapping of >90%). Moreover, we recovered circular extrachromosomal elements, including those linked to antimicrobial resistance, which were also detected in external dog gut datasets. In conclusion, we provide a high-quality reference resource and demonstrate the power of deep long-read metagenomics to resolve microbial diversity in complex host-associated microbiomes.

## INTRODUCTION

Pet dogs (*Canis lupus familiaris*) are commonly regarded as family members, living in close contact with their humans and often sharing not only physical spaces but also microbes^1,2^. Understanding dog microbiome provides insights not only directly linked to their health^3^, but also to household health^4,5^ and global health^6^, being key within One Health frameworks^2^.

Studies on dog gut metagenomics are scarce and all comprehensive ones are based on short-read sequencing^7–12^. Recently, a large-scale project significantly expanded the dog gut metagenomes available^12^. However, most short-read derived metagenome-assembled genomes (MAGs) are heavily fragmented and fail to assemble ribosomal genes. Although the Minimum Information about a Metagenome-Assembled Genome (MIMAG) criteria for high-quality include recovering these (and other elements)^13^, most short-read based studies consider only (estimated) completeness and contamination. We will follow previous literature^14^ and use the term near-finished for MAGs that fulfill all MIMAG high-quality criteria. For instance, in the Unified Human Gastrointestinal Genome catalog, only 12% of the species-level representative metagenome-assembled genomes (573 out of 4,644 MAGs) met these quality criteria^15^. Antibiotic resistance genes (ARGs), mobile genetic elements, and extrachromosomal elements are similarly often absent from short-read MAGs^16–18^. Long-read technologies can address these limitations^19–23^, but large cohorts with deeply sequenced samples using long reads are still scarce. For example, the dog gut microbiome using long-read metagenomics has been recently explored in three pilot studies, which included a single animal^24–26^.

Finally, most large dog gut microbiome studies characterize animals living in colonies, usually from nutritional companies^7,9,12^ rather than pet dogs^10,11^. These cohorts comprise dogs with homogeneous characteristics, such as a limited number of breeds, individuals of similar ages, similar diets, and a shared environment. While these controlled conditions are ideal for certain types of studies (*e.g.,* randomized controlled trials) as they reduce variability, they may not generalize to pet dogs living in households^27^.

Here, we aim to expand the dog gut microbiome data by using deep metagenomics sequencing (minimum per sample: 20 Gbp of long reads + 20 Gbp of short reads) from a relatively large and well-characterized cohort of 51 pet dogs living in an urban environment (Shanghai, China). We make available ∼1,600 Gbp of Nanopore data and ∼1,000 Gbp of Illumina data, 2,676 canine MAGs, 185 circular extrachromosomal elements, non-redundant gene and smORFs catalogs, as well as associated functional and ARGs annotation. By using external public datasets and a new cohort of pet dogs from Germany (also available), we demonstrate that the Shanghai MAG catalog captures global pet dog diversity.

## RESULTS

### Near-finished MAGs are recovered from deep long- and short-read sequencing of a Shanghai dog cohort

We collected a fresh fecal sample from 51 Shanghai pet dogs, associated with extensive dog-associated information (gathered through a questionnaire). Each fecal sample was deeply sequenced using a minimum of 20 Gbp of long-read (Oxford Nanopore Technologies) and 20 Gbp of short-read (Illumina) DNA sequencing (Figure 1A, Supplementary Figure S1).

**Figure 1.**
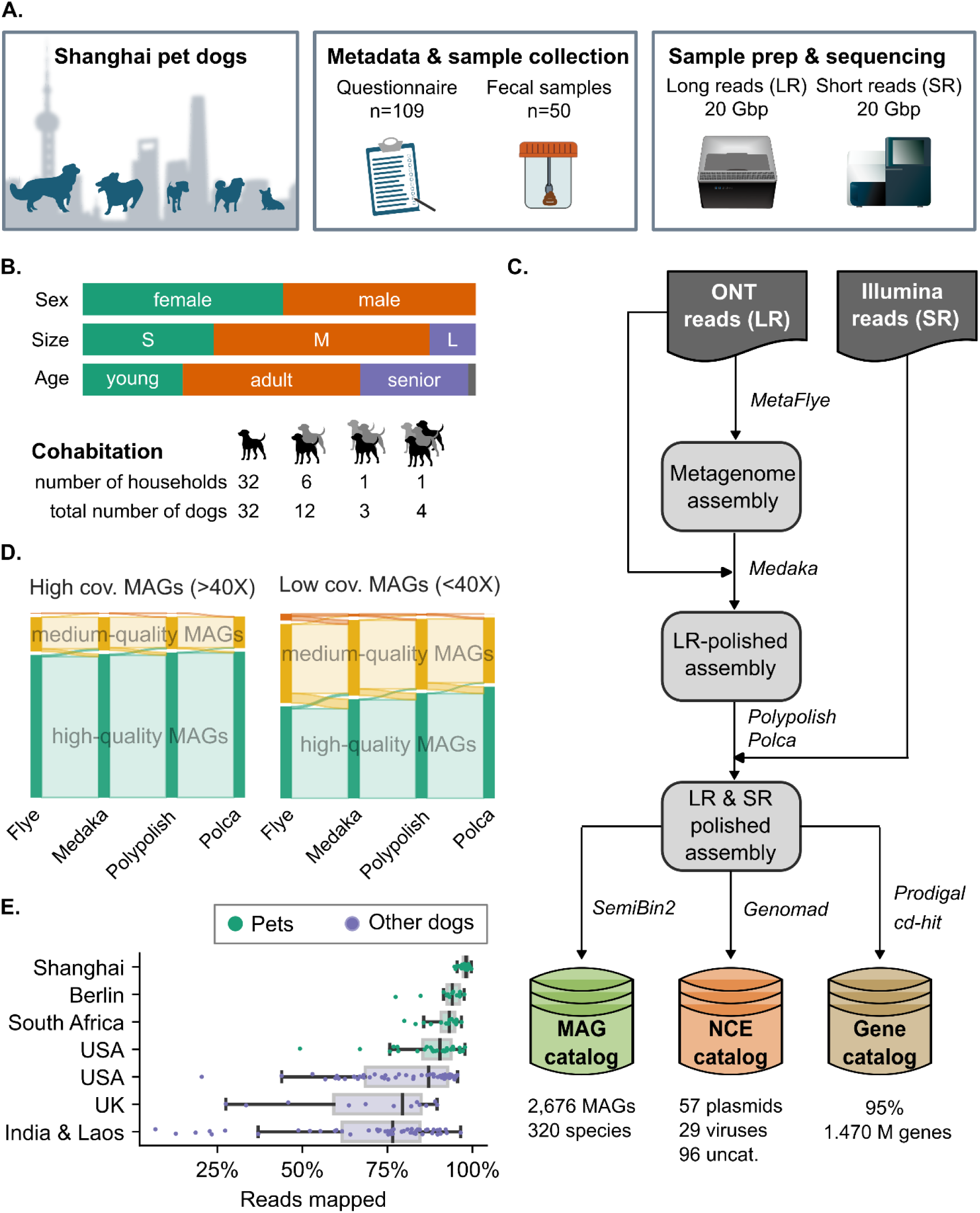
Shanghai pet dog gut microbiome using long-read metagenomics. **A)** Project design overview: we collected information from 109 pet dogs in Shanghai and 51 fecal samples. The samples were processed and sequenced to obtain ≥20 Gbp of both long- and short-read data. **B)** Overview of the Shanghai dog-associated metadata. Complete information for all the dogs can be found in Supplementary Table S1. **C)** MAG generation pipeline: long-read (LR) sequencing data were used for metagenome assembly with MetaFlye. The metagenome assembly was first polished with long reads using Medaka, and later with short reads (SR) using Polypolish, followed by Polca. The polished metagenome assembly was binned using SemiBin2 to obtain the Shanghai Dog MAG catalog (the binning strategy is detailed in the Methods and represented in Supplementary Figure S2). **D)** Short-reads (SR) for polishing increased the quality of the MAGs, especially for those with low coverage (<40X). **E)** The Shanghai dog MAG catalog captures the vast majority of the microbial diversity of other pet dog cohorts living in households (median read mapping of >90%). The mapping is lower for non-pet cohorts (colony, shelter, or free-roaming dogs).

The pet dogs were from the same urban environment (Shanghai, China) and had an urban indoor lifestyle. The cohort comprises animals of different ages, sizes, breeds, diets, and overall habits (Figure 1B and Supplementary Table S1). In addition to the sequenced dogs, we make available 59 questionnaire responses from other Shanghai pet dog owners who did not donate a fecal sample (Supplementary Table S2).

In the MAG generation pipeline, we used the long reads for metagenome assembly and the short reads for polishing, followed by a combination of binning strategies and a final dereplication step, leading to 2,676 MAGs (Figure 1C, Supplementary Figure S2). Each polishing step significantly increased the completeness and decreased the contamination of the canine MAGs (Wilcoxon FDR corrected p-value < 0.05). Polishing converted 177 medium-quality MAGs into high-quality ones, and 48 low-quality MAGs into medium-quality ones (Figure 1D, Supplementary Figure S3), with the greatest benefits observed for MAGs with lower sequencing depth (<40X). Therefore, polishing with short reads was still relevant, especially when working with metagenomes that often include low-abundant species.

The final MAG collection consists of 2,676 MAGs from the Shanghai pet dogs, representing 320 bacterial species (Figure 2A, Supplementary Table S3). The Shanghai dog MAG catalog contains ∼72% near-finished MAGs (n=1,928) (high-quality MAGs fulfilling all the MIMAG criteria: >90% completeness, <5% contamination, presence of ribosomal genes, and at least 18 tRNAs^13^). Of those, 34% were single-contig assemblies, whereas the remaining had a median of six contigs per genome assembly. Medium-quality MAGs represented ∼27% of the MAGs in the catalog (n=732), and only ∼1% (n=16) of the MAGs were high-quality MAGs (>90% completeness, <5% contamination) but did not fulfill some of the remaining MIMAG criteria. Apart from MAGs, we also identified 185 non-redundant circular extrachromosomal elements: 58 plasmids, 30 viruses, and 97 uncategorized elements.

**Figure 2.**
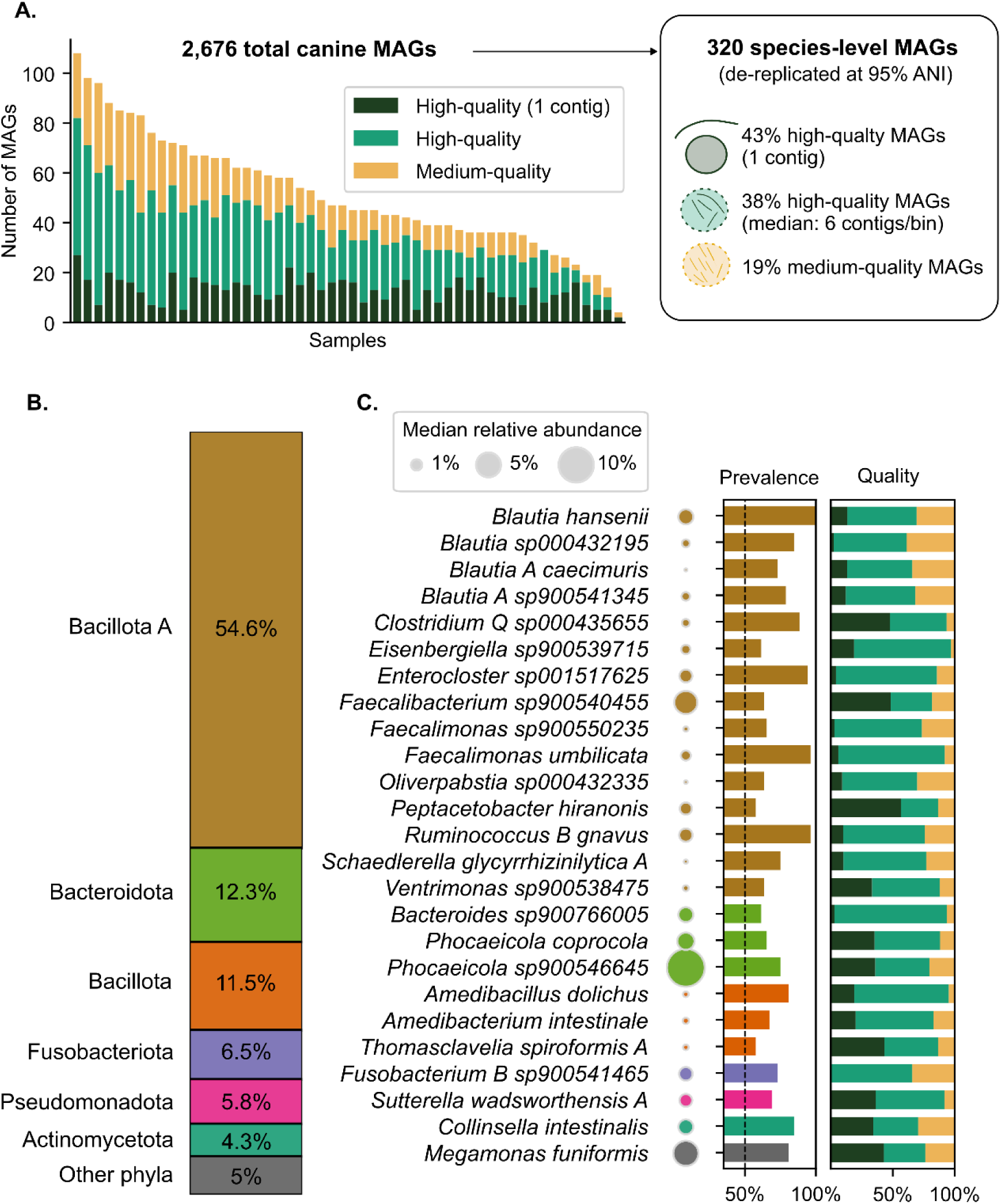
The Shanghai pet dog MAG catalog contains 2,676 MAGs from 320 species. **A)** Stacked bar plots show the total number of MAGs per sample, stratified by quality. Almost all the high-quality MAGs (all but 16) fulfilled the MIMAG criteria. The catalog contains a total of 2,676 MAGs, from which 320 species-level MAGs can be obtained (dereplicated at 95% ANI). **B)** Distribution of phyla in the MAG catalog of Shanghai pet dogs. **C)** The most prevalent species in Shanghai pet dog gut microbiomes, defined as species-level genomes assembled in >30 different dogs (out of 51 total samples). Bubble size represents the median relative abundance of each bacterial species within the total microbiome composition, when present (i.e., zero values were excluded). Prevalence bars show the prevalence for each species. Quality bars show the MAG assembly quality distribution per bacterial species (as defined in **A**). The complete Shanghai Dog MAG catalog information is available in Supplementary Table S3.

Notably, 19 out of the 51 dogs in the Shanghai cohort came from 8 multi-dog households, each contributing at least two sampled dogs (Figure 1B). Within these households, dogs shared a higher proportion of bacterial strains and showed greater similarity in their extrachromosomal elements, as measured by presence-absence data using Jaccard similarity (Supplementary Figure S4).

### The Shanghai dog MAG catalog captures most microbial diversity found in pet dogs

The Shanghai dog MAG catalog captures the microbial diversity of pet dogs living in households globally, as demonstrated by a median read mapping of >90% against metagenome datasets from different geographical origins (Germany, newly sequenced for this study; South Africa^11^; and USA^10^). Despite mapping rates being lower for non-pet dog cohorts, the median was >86% for dogs living in colonies^7,9^ and >75% for dogs living in shelters or the streets (as free-roaming dogs)^11^. Within the Shanghai cohort, 98.2% (median value) of the short-reads mapped to the MAG collection, demonstrating that high-and medium-quality MAGs capture almost all of the diversity present in the samples (Figure 1E, Supplementary Table S4).

More than half of the Shanghai dog MAGs belonged to the Bacillota A phylum, followed by Bacteroidota, Bacillota, Fusobacteriota, Pseudomonadota, and Actinomycetota (Figure 2B, Supplementary Table S3 for the full MAG catalog, annotations with the Genome Taxonomy Database (GTDB) taxonomy r214). Dog gut microbiome studies agree on these phyla being the main ones inhabiting the gut of healthy dogs^3,7^. The most prevalent bacterial species on the Shanghai dog cohort belonged to the Lachnospiraceae family (Bacillota phylum, n=13), followed by the Bacteroidaceae family (Bacteroidota phylum, n=3) (Figure 2C). The most prevalent species in the Shanghai dog cohort–present in almost every dog–were four Lachnospiraceae species: *Blautia hansenii*, *Enterocloster sp001517625*, *Faecalimonas umbilicata*, and *Ruminococcus B gnavus*. Except for *E. sp001517625*, these three species were reported as the most prevalent (prevalence >0.83) on a large cohort of pet dogs living in households in the USA (n=286) as screened with full-length 16S rRNA gene^28^. When focusing on species that are prevalent and also abundant (>2% of total median abundance, when present), we detected *Phocaeicola coprocola*, and *Phocaeicola sp900546645* (Bacteroidaceae family); *Fusobacterium A sp900555845*; *Megamonas funiformis*; and *Faecalibacterium sp900540455* (Figure 2C).

### Species-level long-read MAGs capture novel species and improve representative genome assemblies

Two-thirds of the species-level MAGs represented known bacterial species (∼68%, n=218), as their genomes are found in the GTDB database (r214) (Figure 3A). We compared the species-level high-quality MAG assemblies here to their specific high-quality representative genome in the public database (156 species with either GenBank or RefSeq assemblies) (Figure 3C and Supplementary Table S5). We found that the species-level long-read MAGs were significantly more contiguous, contained more full-length ribosomal genes and unique tRNA genes (all pairwise comparisons had a corrected p-value < 0.05, Supplementary Table S5). They usually fulfilled the MIMAG criteria, which was not true for most of the high-quality representative genomes in public databases. One challenge for short-read shotgun metagenomics is the assembly of ribosomal genes, which are multi-copy and conserved within similar genomes, and consequently, they end up collapsed and unassembled^16^.

**Figure 3.**
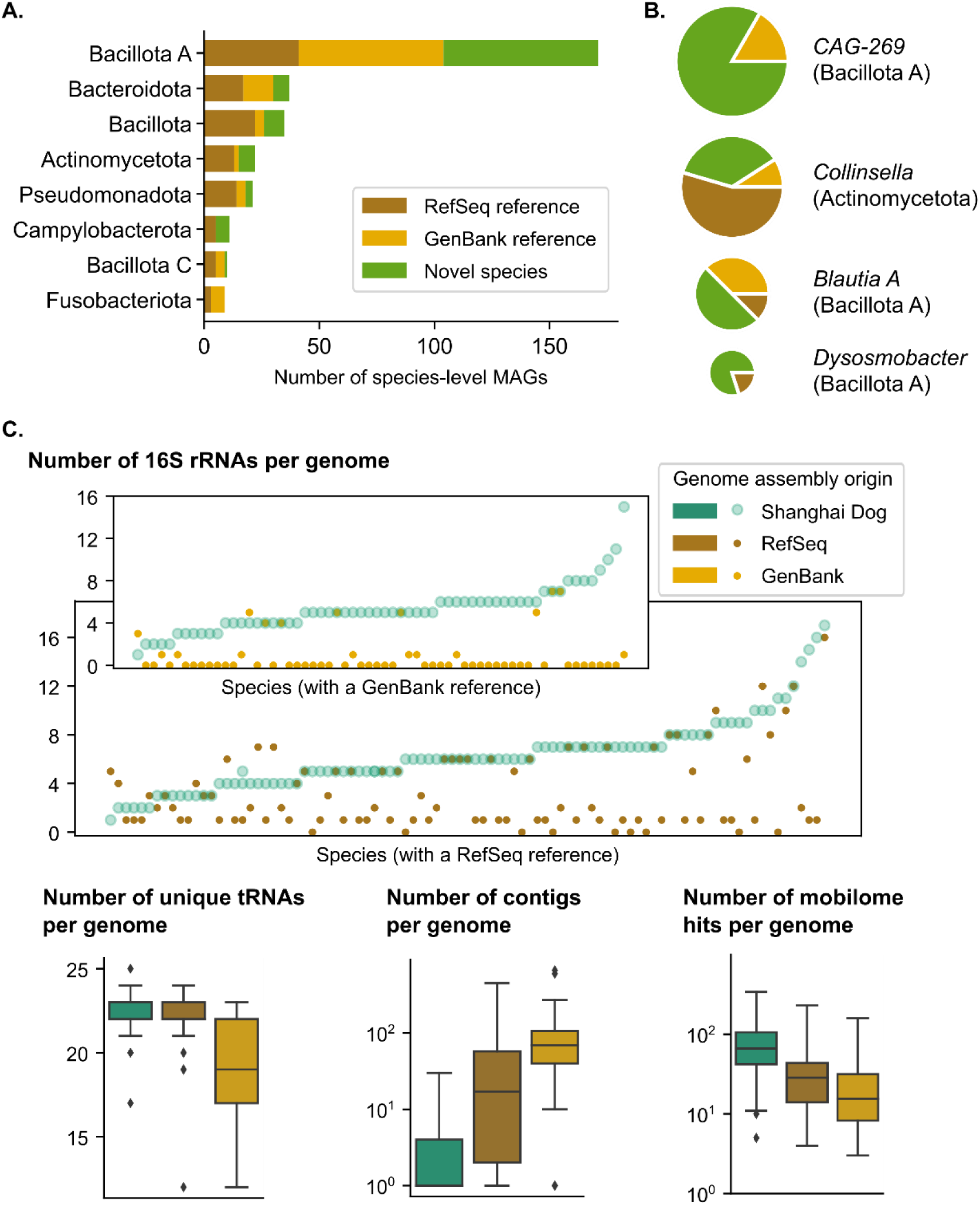
Species-level canine MAGs characterization. **A)** Counts of species-level representative MAGs by phylum in the Shanghai Dog cohort. Stacked bars are colored according to the genome assembly novelty for that species: novel genome (light green); representative species genome assembly present in RefSeq (brown); or representative species genome assembly present in GenBank (yellow). **B)** Top four genera with a larger number of novel species-level MAGs in the Shanghai dog cohort. Pie charts represent the number of species-level genome assemblies within each genus. The size of the pie charts represents the total number of species per genus. **C)** Considering only high-quality genome assemblies (>90% completeness and <5% contamination criteria), comparison of the representative species-level genome assemblies: canine long-read MAGs *vs.* public database representative (RefSeq or GenBank). Scatterplot distribution of the number of 16S rRNA genes: on the x-axis, we represented all the species, and on the y-axis, the number of rRNA genes. Boxplots represent the distribution of the number of unique tRNA genes, number of contigs, and number of mobilome hits, stratified by the origin of the corresponding publicly available representative genome assembly for each species.

Moreover, compared to reference assemblies, canine long-read MAGs usually had a larger mobilome fraction—genes annotated as COG category X: Mobilome (corrected p-value < 0.05, Supplementary Table S5), consistent with previous reports showing that short-read assemblies were largely ineffective for mobile genetic elements analysis^17^, whereas long-reads enabled the capture of mobile elements within their genomic context^19,29^. Overall, our long-read MAGs improve representative genome assemblies on databases, especially those from short-read based assemblies.

One-third of the species-level MAGs represented bacterial species not present in the GTDB (r214) (∼32%, n=102). The genera that contained the most novel species were *CAG-269* with 10 different new species-level MAGs, followed by *Dysosmobacter*, *Collinsella*, and *Blautia A* with four different new species-level MAGs each (Figure 3B, Supplementary Table S3). To further explore the novelty of the species and considering that 16S rRNA gene databases are more extensive than whole genomes, we aligned the full-length 16S rRNA genes of the high-quality MAGs against the 16S ribosomal RNA sequences database from NCBI using the blastn suite. The best hits corresponded to 16S rRNA genes from known species within the same genera, and in 11 cases, the identity of the best hit was >99% (Supplementary Table S6). In nine out of 11 cases, we confirmed the inability of this marker gene to distinguish species within certain genera^30^.

Several bacterial species lack representative genome assemblies in public databases but have phenotypic and 16S rRNA gene sequencing data. These include *Sutterella stercoricanis* and *Gluceribacter canis*, which were first isolated in dog feces^31,32^. A previous study showed that high-quality long-read MAGs can link 16S rRNA gene-based community profiling studies to the functional potential of the genomes within the same environment^20^. Long-read MAGs enabled us to putatively link genome assemblies to previously phenotypically characterized species in dogs through 16S rRNA gene sequences.

### Long-read metagenomics uncovers ARG-carrying circular extrachromosomal elements in the canine gut

We identified 253 unique antibiotic resistance genes (ARGs) in the Shanghai dog cohort (using RGI, see Methods). Tetracycline ARGs were the most common, representing ∼32% of the total ARG counts (with 2,461 total hits, corresponding to 25 unique ARGs), followed by aminoglycoside and lincosamide ARGs, representing ∼13% of the total ARG counts each (1,021 and 1,001 total hits, corresponding to 31 and 5 unique ARGs, respectively). This aligns with findings from a large-scale resistome analysis including >3,000 dog samples, where tetracyclines, lincosamide, and aminoglycosides ARGs were the most abundant ARGs on dogs^33^. Additionally, these ARGs correspond to antimicrobial drug classes that are essential and commonly used in small animal medicine^34^.

Interestingly, of the 253 unique ARGs, 43 were exclusively found in contigs that were not part of a MAG. Among ARGs, those located on extrachromosomal elements (ECEs) are particularly concerning due to their potential for horizontal gene transfer across bacterial species. This mobility facilitates rapid dissemination of antimicrobial resistance within microbial communities^35^. The use of long-read sequencing is not only crucial for contextualizing ARGs, but also effective at assembling circular ECEs that might harbour them^19,21^. In addition to the MAG catalog, we assembled 185 non-redundant circular ECEs: 58 plasmids, 30 viruses, and 97 uncategorized elements. Among those, four plasmids and six uncategorized elements harboured ARGs (Figure 4, Supplementary Table S7).

**Figure 4.**
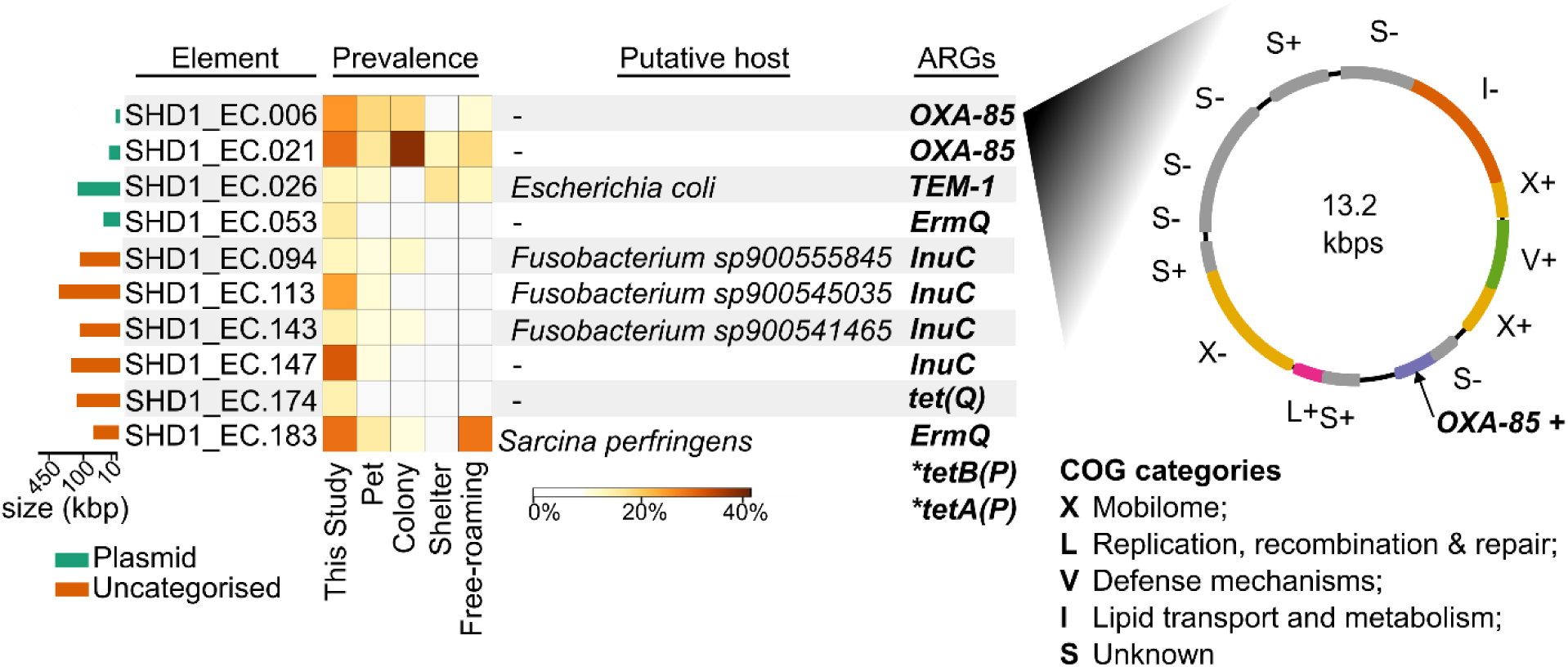
Circular extra-chromosomal elements (ECEs) harboring antimicrobial resistance genes (ARGs) in the Shanghai Dog cohort. The heatmap represents the prevalence of each ECE in both our cohort and external dog cohorts (as confirmed by read mapping). For some ECEs, we predicted their putative bacterial host and identified their ARGs. The circular diagram represents the ORFs identified in EC.006 and their predicted COG functional category.

We identified two different, circular, and single-contig ECEs harboring the *OXA-85* gene, which were present in more than 25% of the samples in our cohort, and had a prevalence of up to 45% in colony dogs (SHD1_EC.021) and up to 14% in other pet dog cohorts (SHD1_EC.006) (Figure 4). The *OXA-85* gene encodes a class D β-lactamase that confers narrow-spectrum resistance primarily to penicillin-class antibiotics^36^. We could not assign these two ECEs to a putative bacterial host; therefore, they could be present in multiple species. Previously, *OXA-85* had only been reported on the chromosome of *Fusobacterium nucleatum*^36^ and *Fusobacterium necrophorum*^37^, consistent with entries in the CARD database^38^, which additionally found it in *Campylobacter ureolyticus*. In fact, we also detected *OXA-85* within the chromosomes of *Fusobacterium B sp900541465* (n=3), *Fusobacterium A sp900543175* (n=1), as well as *Phascolarctobacterium A sp900552855* (n=3). This finding is concerning due to the extrachromosomal location of the *OXA-85* gene, which could enable horizontal gene transfer; its potential association with unreported bacterial hosts; and its prevalence across diverse dog cohorts. All these suggest broader dissemination and host range than previously observed.

We found one circular ECE harboring *TEM-1*, which was linked to *E. coli* as its putative host. *TEM-1* is a plasmid-encoded beta-lactamase–found in many Gram-negative bacteria–that confers resistance to penicillins and first-generation cephalosporins^39^. Its prevalence across other dog cohorts reinforces concerns about the circulation of clinically relevant ARGs in non-clinical reservoirs. We also identified two different, circular, and single-contig ECEs carrying the *ErmQ* gene, one with *Sarcina perfringens* as its putative host (SHD1_EC.183) –consistent with the original *ErmQ* report^40^. The *ErmQ* gene encodes a ribosomal RNA methyltransferase that confers resistance to macrolide, lincosamide, and streptogramin B antibiotics. Interestingly, a recent paper characterizing pet-derived *S. perfringens* in China found that 91% of them showed a high resistance rate to erythromycin associated with the presence of the *ErmQ* gene^41^. Finally, and coinciding with the most abundant and prevalent ARGs, lincosamide (*lnuC*) and tetracycline resistance genes (*TetQ*, *TetA*, *TetB*) were the ARGs detected in the remaining five ECEs (Figure 4).

### Global analysis detected that dogs living in colonies present a different gut microbiome

To contextualize our data and investigate factors that structure the dog gut microbiome, we integrated the Shanghai dog gut microbiome with a set of external dog samples (Berlin cohort, and publicly available canid cohorts until February 2023). The largest dog gut metagenome studies focused on colony dogs rather than pet dogs living in households^7,9^. Only three studies involved pet dogs: one examined dogs with recurrent diarrhea^8^, while the other two analyzed healthy pets from South Africa^11^ and the USA^10^. Using representative samples from these public datasets (Supplementary Table S8), we assessed the influence of the living environment (household, colony, or free roaming), age, size, and sex of the animal. Since these studies are short-read studies, for consistency, we used the short-read Illumina component of our data and performed taxonomic profiling on the combined dataset using a pipeline based on singleM^42^ (see Methods).

The living environment was the variable structuring the microbial community taxonomic composition at the beta diversity level (PERMANOVA adjusted p-value = 1.67 x 10^-3^), with a clearly separated cluster for dogs living in colonies. Pet dogs overlapped and clustered in the ordination plot, even though they were from different geographies (China, USA, Germany, and South Africa) (Figure 5B). Analogously, alpha diversity (Shannon index) was significantly higher in colony dogs when compared to pet dogs (Mann-Whitney U corrected p-value = 4.90 x 10^-3^) and wild canids (Mann-Whitney U corrected p-value = 0.025; Figure 5A). Ancient coprolite samples presented the lowest alpha diversity values and clustered independently on the beta diversity plot, but considering that these samples all originate from the same study^43^, it was not possible to exclude a study effect (Supplementary Figure S5).

**Figure 5.**
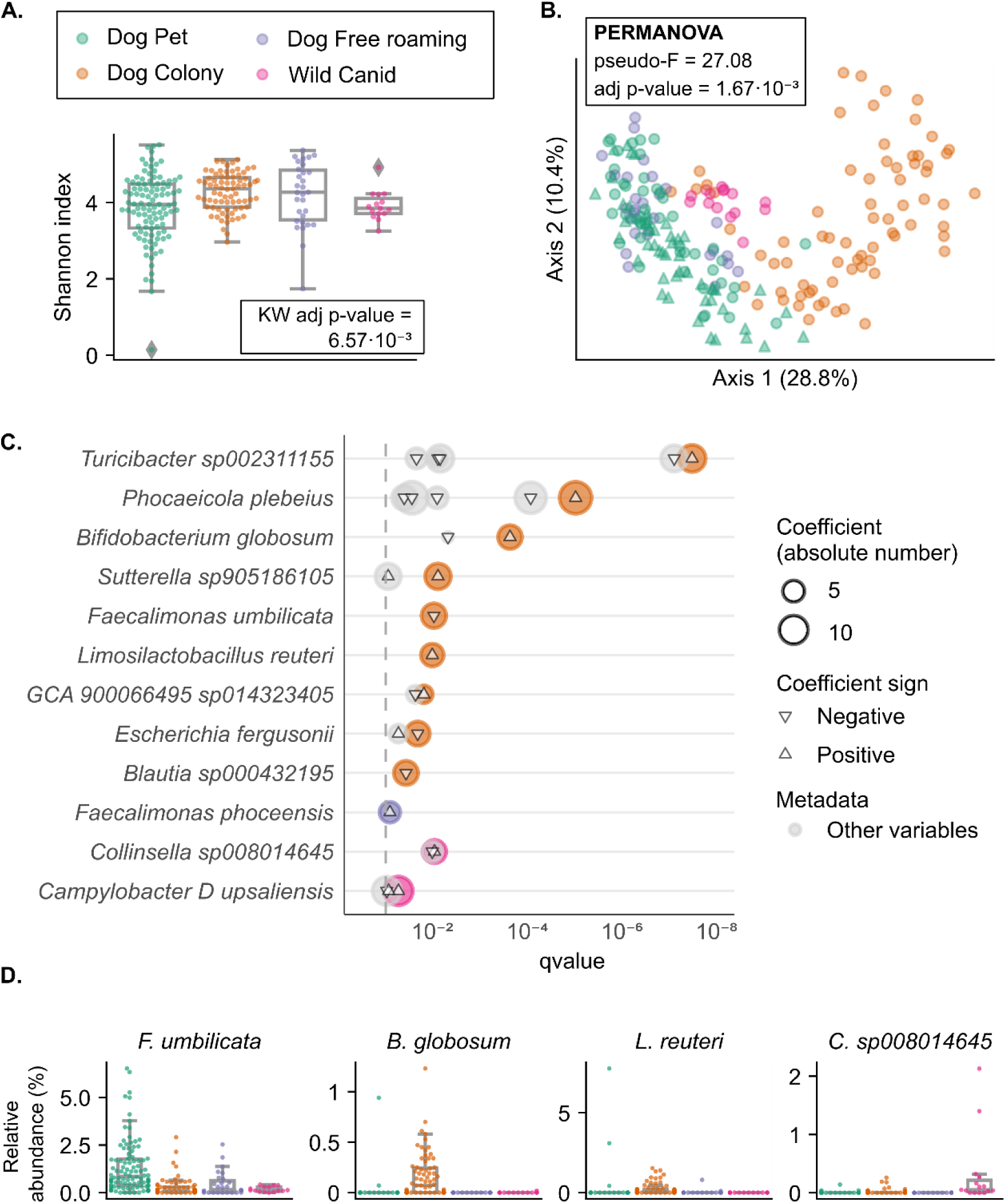
The living environment is the major determinant of dog gut microbiome composition. The colored legend applies to the whole figure. **A)** Boxplots representing alpha diversity (Shannon index). **B)** PCoA plot representing beta diversity (Bray-Curtis on log-transformed data). Green triangles indicate pet dogs in this study. **C)** Differentially abundant bacterial species in each dog group when compared to pet dogs (Maaslin2 and Kruskal-Wallis, see Methods). The x-axis shows the q-value of each species computed by Maaslin2. Circles are colored according to the group to which the dogs belong. Grey circles represent significant differences in that species, for variables other than the living environment. Diversity analysis with all dog groups can be found in Supplementary Figure S5. **D)** Boxplots of total relative abundances (%) of selected differentially abundant bacterial species, considering the living environment.

When looking at differential species abundance, six species were more abundant in colony dogs compared to pet dogs and other dog cohorts, including *Bifidobacterium globosum* and *Limosilactobacillus reuteri* (Figure 5C and 5D). *Lactobacillus* and *Bifidobacterium* species are commonly regarded as probiotics in dogs^44^, so this could represent a more common intake of probiotics in dogs living in colonies compared to other dog cohorts. Moreover, we identified *Faecalimonas umbilicata* as a pet-associated species, as it was significantly more abundant in pet dogs compared to all the other dog groups. Finally, we also detected two bacterial species more abundant in wild canids and one in free-roaming dogs (Figure 5C).

Differential abundance analysis also detected bacterial species associated with: age of the animal (n=13), body condition (n=2), and size of the animal (n=2) (Supplementary Figure S6). However, no significant differences were found regarding sex. Differences related to dogs’ age were mostly significant when comparing senior to young animals, *Dorea B phocaeensis*, *Slackia A piriformis*, *Eisenbergiella sp900539715,* and *Faecalibacterium sp900540455* being the most significant. Interestingly, *Faecalibacterium* and *Slackia* were among those genera more abundant in healthy dogs when compared to those having cancer^45^, and cancer is an age-related disease. We also found *Butyricicoccus pullicaecorum* to be more abundant in young dogs. Interestingly, this species has been assessed as a potential probiotic and has been shown to extend lifespan in a *C. elegans* model^46^.

### Generated resources description

In this study, we generated several high-quality resources from pet dogs using long-read metagenomics, polished with short reads. We make these datasets available to be further explored and used:

1. **Comprehensive dog-associated information of Shanghai pet dogs**: includes responses from 107 dog owners in Shanghai regarding their pets’ health, diet, and lifestyle (among others) collected through an extensive questionnaire (Supplementary Table S2). Considering the most relevant variables, we created a simplified table for the 51 sequenced dogs included in the study (Supplementary Table S1).
2. **Deeply sequenced samples (raw data)**: includes 52 fecal samples belonging to 51 dogs (1 biological replicate) with at least 20 Gbp of data for long-read and 20 Gbp of data for short-read. Long-read datasets have up to 65 Gbp of data per sample (available on ENA under Bioproject ID: PRJEB85799).
3. **Metagenome-assembled genomes (MAG) catalog**: includes 2,676 MAGs that belong to 320 different bacterial species. 72% are near-finished MAGs, meaning they are >90% complete, <5% contaminated, harbor ribosomal genes, and at least 18 unique tRNAs. 665 of the MAGs are single-contig. 32% of the species-level representatives are novel (Supplementary Table S3). In addition to the sequences (in FASTA format), we provide extracted rRNA genes (with the 16S rRNA linked to MicrobeAtlas to enable users to link to 16S amplicon-based studies^47^), and annotations using eggnog-mapper and RGI.
4. **Extrachromosomal elements (ECEs) catalog**: includes 185 single-contig circular elements (non-redundant), as assembled by MetaFlye from our cohort. We identified 58 plasmids, with sizes ranging from 4,333 bp to 154,111 bp (median length: 30,562 bp); 30 viruses; and 97 uncategorized elements. Moreover, we predicted the putative bacterial host for 28% of them (n=52).
5. **Gene catalog**: includes 1,470,802 non-redundant genes (95% nucleotide identity, representing species-level unigenes^48^). Of these, 98.7% are complete genes, and 73.5% are contained in MAGs or ECEs.
6. **Small ORFs catalog**: includes 403,491 non-redundant smORFs, which produce 273,065 clusters at 90% identity.
7. **External dog datasets**: raw data for a newly sequenced short-read metagenomics pet dog cohort from Berlin (Bioproject ID: PRJNA881055), as well as standardized and manually curated metadata for dog gut metagenomics datasets in public databases up to February 2023^7–11,43,49–58^ (Supplementary Table S9). We also created a metadata subset including a single representative sample per animal for longitudinal or repeated measures studies (Supplementary Table S8).

We created a website (https://sh-dog-mags.big-data-biology.org/) to dynamically explore the MAG catalog, and all the derived resources can be downloaded on Zenodo: https://doi.org/10.5281/zenodo.16356977

## DISCUSSION

Here, we present the Shanghai dog gut microbiome resource, comprising samples from 51 pet dogs. This dataset includes extensive metadata –over 50 lifestyle, health, and dietary variables collected through owner questionnaires– and deep metagenomics sequencing. From this, we derived multiple resources, including a comprehensive pet dog MAG catalog with 2,676 MAGs from 320 different bacterial species. Our Shanghai dog MAG catalog captures most of the global microbial diversity of pet dogs as highly contiguous, near-finished MAGs, as shown by the high mapping rates to the catalog (>90% median) of external pet dog cohorts from different geographies (Germany, South Africa^11^, USA^10^).

Previous short-read metagenomics studies on the dog gut have focused on functional and taxonomic profiling^8–11^ or building a gene catalog^7^, without assembling MAGs. Recently, a large-scale study on companion animals retrieved 7,275 high-quality and 21,706 medium-quality MAGs from canine and feline samples using short-read metagenomics^12^. However, short-read metagenome assemblies typically fail to recover ribosomal RNA genes^16^. They also often miss other important features such as antimicrobial resistance genes (ARGs)^18^ and mobile genetic elements^17^. In this study, we present a high-quality canine MAG catalog built from long-read nanopore sequencing, polished by short reads. Most (∼72%) of the MAGs are near-finished, with completeness >90%, contamination <5%, and include ribosomal RNA genes and tRNAs, thus fulfilling the full MIMAG criteria^13^. Moreover, we retrieved 185 non-redundant circular extrachromosomal elements; 1,470,802 non-redundant unigenes; and 403,491 non-redundant small open reading frames (smORFs). Our resource includes 320 species, and over 80% have a near-finished MAG as a representative genome. Many of our species-level representative MAGs improve existing reference genomes in the GTDB r214 database (including ones that were obtained from cultures, but assembled with short reads), offering higher contiguity, and including the ribosomal genes and tRNAs. Looking forward, we expect that many species in multiple habitats will have higher-quality genomes available in the next few years as long-read metagenomics becomes widely used.

A previous study suggested that >40X coverage with Nanopore R10.4 chemistry can yield near-finished MAGs from metagenomes^14^. Here, we observed further improvements in completeness and contamination values when incorporating short reads, suggesting additive benefits. While we cannot rule out differences from basecalling models—our data were basecalled using the high-accuracy model versus the super-accuracy model used in that study—particularly considering that complex microbiomes contain a plethora of low abundance genomes, it is evident that long-read sequencing is key to generating near-finished MAGs, and, with current ONT chemistry, short read sequencing is still valuable.

Within a One Health framework–which recognizes the interconnectedness of global human, animal, and environmental health–the close interaction between pets and humans, particularly in shared household environments, creates ideal conditions for the bidirectional transmission of bacteria and their associated ARGs^59^. Although we did not target the human gut microbiome, we observed that dogs living in the same household shared significantly more bacterial strains and extrachromosomal elements (ECEs) compared to dogs from different households, confirming the existence of intra-household transmission patterns.

Notably, 10 of the assembled ECEs carried ARGs, which play a central role in horizontal gene transfer and the dissemination of resistance within shared environments^35^. Among these, we identified a putative *E. coli* plasmid carrying the *TEM-1* gene, underscoring concerns about plasmid-mediated antibiotic resistance in a bacterial species that is commonly shared between humans and animals, and known for its ability to mediate gene exchange across mammalian hosts^60,61^. Additionally, the detection of the *OXA-85* gene in two ECEs and across multiple anaerobic species (e.g., *Fusobacterium*, *Phascolarctobacterium*) suggests a mobilization event and an expanded host range for this ARG. Although these specific ARGs are not currently considered major clinical threats, their detection in mobile genetic elements within the pet dog gut microbiome highlights the value of metagenomics surveillance using long-reads to identify ARG reservoirs and monitor ARG flow across taxa and environments^62^.

We observed that the mapping rate to our catalog for samples of dogs living in colonies was slightly lower than pets, which hinted at a difference in their microbiome structure. Previous studies showed that factors such as industrialization and feralization influence the gut microbiome composition and function in companion animals like dogs^11^ and cats^63^. However, in a study by Yargaladaa and collaborators, industrialization was confounded with geography, as each industrialization level corresponded to a distinct geographic region. In contrast, a large-scale consumer study including 192 US pet dogs (16S rRNA gene sequencing) did not detect effects of urbanization on the gut microbiome, though some geographic signatures were reported^64^. The largest companion animal microbiome study to date^12^ reported that housing facilities (research facilities *vs.* household *vs.* strays) significantly influenced the dog gut microbiome. Beta diversity analysis confirmed that pet dogs from different geographic regions (China, Germany, US^10^, and South Africa^11^) presented similar gut microbiome compositions, clustering together and overlapping within the same region of the PCoA plot. In contrast, colony dogs from nutritional research facilities formed a distinct cohesive cluster despite their different geographic origins (US^7^ and UK^9^). Taking all these observations together, we conclude that the living environment has a stronger influence than geography on the composition and structure of the canine gut microbiome. These living environment differences likely reflect differences in multiple factors such as diet, environmental exposure, human interaction, and veterinary care. Thus, while colonies have definite benefits for controlled studies, their microbiome differs from the general pet population, and results from colony studies should be complemented with studies in pets.

In summary, our study significantly expands the existing resources on the dog gut microbiome by generating and sharing a comprehensive multi-terabyte dataset derived from deep long- and short-read metagenomic sequencing of a well-defined cohort of urban pet dogs. With over 2,600 canine MAGs, hundreds of circular elements, and extensive gene and smORF catalogs, this dataset provides an unprecedented foundation for future research. Moreover, by validating our Shanghai-derived MAG catalog against both public data and an independent German cohort, we show that it captures a broad spectrum of global pet dog microbiome diversity, establishing it as a valuable reference for comparative and functional microbiome studies.

## STAR METHODS

### DATA AVAILABILITY

Raw reads and MAG catalog for the Shanghai pet dog cohort were deposited at the European Nucleotide Archive under Bioproject ID PRJEB85799. Berlin pet dog raw data can be found under Bioproject ID PRJNA881055.

MAG annotation files, ARGs prediction, ECEs catalog, gene catalog, and smORFs catalog can be found at Zenodo: https://doi.org/10.5281/zenodo.16356977. Finally, all original code is publicly available at GitHub: https://github.com/BigDataBiology/ShanghaiDogs

### EXPERIMENTAL MODEL AND STUDY PARTICIPANT DETAILS

In this study, we analyzed the fecal microbiome of pet dogs living in households in Shanghai. The Animal Welfare and Ethics Group (Department of Experimental Animal Science, Fudan University) approved this study under the approval reference 022JSISTBI-003. All the dog owners who donated a fecal sample provided informed consent to participate in the study.

### METHOD DETAILS

#### Dog-associated metadata collection

We designed a comprehensive questionnaire that covers various aspects of dogs, including their basic information (breed, age, sex, weight) and other relevant variables related to lifestyle, habits, diet, health, and more. Some of the questions in our questionnaire were adapted from Lehtimäki et al., 2018^65^. The questionnaire was entered into and administered through the *wjx* mini app on WeChat and shared across relevant WeChat groups. 107 dog owners answered the questionnaire, and 51 agreed to donate a fecal sample from their dogs (Supplementary Table S2).

For the 51 dogs sampled, updates in the metadata, such as health status, were done on the sampling date and adapted for further analysis (Supplementary Table S1). We grouped the dogs into three age categories, as previously considered^66^: Young (< 2 years old), Adult (2-6 years old), and Senior (≥ 7 years old). Finally, we classified the dogs according to size, as previously described^67^: I) in five clusters, and II) in size categories: Small (< 10 Kg), Medium (≥ 10 to < 25 Kg), and Large (≥ 25 Kg). The dog cohort was balanced by sex.

#### Fecal sample collection

The owners collected the fecal samples when walking their dogs per their regular habits, which were given to us 0h-24h after the dog’s deposition. Therefore, we have neither altered nor manipulated the animals in any way.

Once we had the fresh dog fecal sample, we used a fresh pair of gloves to cut the feces with a plastic spatula and expose the inner part. We collected a spoon-sized sample from the inner part of the feces using a tube with a spoon. We did this process twice per sample, so we have a biological duplicate as a backup. Immediately after processing, the tubes were stored in dry ice and transported to a −80 °C freezer until further processing.

#### DNA extraction and sequencing

Fecal samples were processed and sequenced by Novogene (Beijing, China). The DNA was extracted using the CTAB method. Genomic DNA quality was verified by: 1% agarose gels to detect DNA degradation; Nanodrop OD 260/280 ratio to check the purity of DNA; and Qubit® DNA Assay Kit (in Qubit® 3.0 Fluorometer) to measure DNA concentration.

For short-read Illumina sequencing library preparation, 0.2 μg of DNA per sample was used. The DNA was sonicated to an average fragment size of 350 bp. It was generated using NEB Next® Ultra™ DNA Library Prep Kit for Illumina (NEB, USA) following the manufacturer’s recommendations, and index codes were added to each sample. Clustering of the index-coded samples was performed on a cBot Cluster Generation System using the Illumina PE Cluster Kit (Illumina, USA), following the manufacturer’s instructions. The final DNA libraries were sequenced on an Illumina NovaSeq 6000, generating 150 bp paired-end reads.

For long-read Nanopore sequencing library preparation, 1-2 μg of DNA per sample was used. The DNA was size selected using magnetic beads, and the library was constructed following the Oxford Nanopore Technologies (ONT) 1D Genomic DNA by Ligation protocol (v14) with the Ligation Sequencing Kit SQK-LSK114 following the manufacturer’s recommendations. The final DNA libraries were sequenced on a PromethION P48 instrument using R10.4.1 flow cells. Raw signal data (fast5 files) were basecalled using Guppy v6.4.6 with the High-Accuracy (HAC) model.

#### Selection of external dog gut metagenomes

We selected all the available dog gut metagenomes until February 2023. We used the GMGC catalog^48^ to collect the initial entries and their associated metadata, and we added new studies by searching at PubMed and the SRA database. Dog-associated metadata was manually curated from each sample’s associated literature and Biosample or SRA-associated metadata. We additionally included a new shotgun metagenomic dataset of 15 pet dogs from Berlin (Germany).

The collection of external dog metagenomes used in this study and their associated metadata can be found in the Supplementary Table S9. Whenever multiple samples from the same animal were available, we selected a single representative sample (Supplementary Table S8). For choosing the representative samples, we prioritized that the dogs were on a non-interventional ‘baseline’ diet, the healthiest status (*e.g.,* absence of clinical signs after treatment, in cases of chronic enteropathies), or the sample with the highest throughput. Samples that could not be linked to their metadata were excluded from the representative set of samples.

In addition to publicly available cohorts, we included a newly sequenced cohort from Berlin with 15 pet dogs from households in the context of a food allergy and tolerance study^68^.

#### Reads pre-processing

Before the read pre-processing step, we run MinIONQC^69^ on ONT raw reads to evaluate the quality of the sequencing runs, including the q-score, sequence length, and total throughput.

Short and long reads were pre-processed by quality, length, and presence of host reads. For Illumina short reads, we used NGLess^70^, and raw reads were discarded if they presented a quality score lower than 25, were less than 45 bp in length, or mapped to the dog genome. For Nanopore long reads, we used Chopper^71^, and raw reads were discarded if they had a quality score lower than 10, were less than 500 bp in length, or mapped to dog genomes. After Chopper, we ran Porechop_abi^72^ to remove any remaining adapters and discard reads with middle adapters.

#### Metagenome assembly and binning

Pre-processed long reads were used to perform metagenomics assembly with MetaFlye^73^. We performed two to three metagenome assemblies per sample, depending on the initial sequencing depth: all data, 20 Gb, and 10 Gb. Metagenomics assemblies using the whole sequencing data (all data) are the norm throughout the manuscript. Subset metagenomics assemblies were only used to obtain MAGs representing species that were missed with the ‘all data’ strategy (Supplementary Figure S2). The data subsets were generated with Rasusa^74^, indicating the desired sequencing depth.

The MetaFlye metagenome assemblies were further polished using three consecutive polishing steps: Medaka, which used long reads for polishing; followed by Polypolish^75^, which used short reads for polishing; and a final round using Polca^76^, which also used short reads for polishing. After polishing, we binned the contigs using SemiBin2^77^ with three different strategies for ‘all data’ assemblies: single-sample binning with short reads, single-sample binning with long reads, and multi-against-multi binning^78^. For the subset assemblies, we exclusively used single-sample binning with short reads.

#### MAG catalog generation and characterization

To characterize the generated metagenome bins, we used CheckM2^79^ to evaluate contamination and completeness, followed by GTDB-tk^80^ with the GTDB r214 database to assign taxonomy to the genomic bins. We used DASTool^81^ to dereplicate the metagenome bins originating from the same initial read set (all data) with different binning strategies. Finally, we manually included metagenome bins from the subset assemblies if they represented a different taxonomy not captured with the ‘all data’ assembly.

The final collection of MAGs includes only those metagenome bins meeting high- or medium-quality standards based on the MIMAG criteria^13^ (Supplementary Table S3). According to these criteria, high-quality MAGs have >90% completeness, <5% contamination, and must contain ribosomal genes and at least 18 out of the 20 tRNAs. Medium-quality MAGs are defined by >50% completeness and <10% contamination. To distinguish between MAGs meeting the full MIMAG high-quality criteria and those meeting only completeness and contamination thresholds (>90% completeness, <5% contamination), we refer to the former as *near-finished MAGs* and the latter as high-quality MAGs throughout this study.

Apart from checking completeness and contamination, we used barrnap to identify the ribosomal genes, and for cases where the 5S rRNA genes were not predicted, we also used RNAmmer^82^. The extracted 16S rRNA genes were mapped to MicrobeAtlas OTUs^47^ using their online API. We predicted tRNAs using tRNAscan^83^. Finally, the species-level MAG representatives were obtained by running dRep^84^ at 95% ANI.

#### Manual quality curation on the Shanghai Dog MAG catalog

During the quality assessment of *Allobaculum stercoricanis* MAGs, we observed that high-quality ones were downgraded to medium-quality after polishing. This shift in quality classification coincided with a change in the default model chosen by CheckM2 during its evaluation (moving from a general to a specific model). Several of these MAGs were single-contig, circular genome assemblies –which typically indicate high completeness and low contamination– but still classified as medium-quality. To address this discrepancy, we manually re-evaluated the quality of *A. stercoricanis* MAGs using the general CheckM2 model throughout all the polishing steps.

Four out of the total six *Klebsiella pneumoniae* MAGs identified in our cohort, derived from consecutive canine fecal samples (D023–D026), presented an identical ARGs pattern and exhibited extremely high ANI values. Samples D023–D026 are all from different households and different neighbourhoods of Shanghai; therefore, we suspected a cross-contamination event had happened during the sample processing. Dog D024 was on antibiotics, and its microbiome consisted of a *K. pneumoniae* MAG with 6,000X coverage and some other very low abundant bacteria potentially contaminating nearby samples.

#### Bacterial strain and extrachromosomal elements sharing

We compared the sharing patterns of bacterial strains and extrachromosomal elements in the Shanghai dog cohort, considering pairwise comparisons between dogs within the same households *vs.* between households.

To analyze the sharing patterns of bacterial strains, we divided the number of shared bacterial strains (considering ‘same strain’ those MAGs with ANI > 99%) by the number of shared bacterial species (considering ‘same species’ as those MAGs with ANI > 95%). Two samples were only compared if they presented at least 10 shared bacterial species.

To analyze the sharing patterns of extrachromosomal elements, we used the ‘covered fraction’ table generated by CoverM^85^ –based on short-read mapping results to the MAG and extrachromosomal element catalogs. We converted this table into a presence-absence matrix by considering elements with >80% coverage as present. Considering the extrachromosomal elements present in at least 10% of the samples, we calculated sample similarity using Jaccard Similarity.

To evaluate the statistical significance of shared bacterial strains or extrachromosomal elements between dogs from the same household *vs.* different households, we performed a Mann–Whitney U test.

#### Gene catalog generation

Open reading frames were predicted with Prodigal in metagenomics mode (-p meta)^86^. ORFs were then clustered at 100% nucleotide identity (with 100% coverage of the shorter sequence) by comparing shorter sequences to longer sequences. This catalog was then clustered at 95% nucleotide identity (with 90% coverage of the shorter sequence) using CD-HIT-EST (command line parameters, −c 0.95 −d 0 −g 1 −G 0 −aS 0.9)^87^.

#### smORF catalog generation

We ran GMSC-Mapper 0.1.0^88^ on the polished metagenome assemblies and recovered the non-redundant small ORFs (with 100% amino acid identity) by removing duplicated sequences. These smORFs were subsequently clustered at 90% identity using CD-HIT with parameters −c 0.9 −n 5 −d 0.

#### Functional annotation of MAGs

Using polished metagenome assemblies, we predicted protein-coding genes using Prodigal 2.6.3 with ‘-p meta’ option. Using as input the predicted proteins file, we run eggNOG mapper 2.1.12 with the following options: ‘-m diamond --itype proteins’.

#### Antimicrobial-resistant genes prediction

We predicted the ARGs using two input file types: polished metagenome assemblies (contig-level fasta files) and the MAG catalog. We ran RGI v6.0.3 with the CARD database^38^. We filtered the output table by only keeping those ARGs predicted as ‘strict’ or ‘perfect’ and that presented at least 90% identity and 90% coverage to the hit on the database.

For the website (https://sh-dog-mags.big-data-biology.org/), we only included ‘clinically-relevant’ ARGs, which were obtained by filtering the output ARGs to include those also present in the ResFinder v4.0 database^89^ by using the database mappings from argNorm 1.0^90^.

#### Putative plasmid mining and identification

For all samples, contigs identified by Flye as circular were extracted and RecurM v0.2.8 was run on them with the options ‘-c 2 –collapse_against_assembly’. Only circular clusters identified by RecurM were considered for further analysis. To label the generated circular clusters, GeNomad v1.1.0^91^ was run with default parameters, and any contigs with at least 1 plasmid hallmark gene as defined by GeNomad were considered a plasmid. Putative plasmids were annotated with DRAM v1.5.0^92^ with default settings to determine functional gene encoding. To identify putative viruses, VirSorter2 v2.2.4^93^ was run with the options ‘--include_groups dsDNAphage, NCLDV, RNA, ssDNA, lavidaviridae --high-confidence-only’ and any viral sequence identified as ‘full’ in the final-viral-score.tsv file was labelled as virus. All other sequences were considered ‘non-categorised’. Any contigs that were binned into a MAG were discarded.

To identify putative hosts for plasmids, a co-abundance approach was used. MAG and putative plasmid abundance was calculated by mapping metagenomic reads using CoverM with the trimmed mean option. The trimmed mean of the relative abundances of both MAGs and putative plasmids was used as input into FlashWeave v0.19.2^94^ using the sensitive, heterogeneous mode to construct a co-abundance table. Relationships between a MAG and a circular cluster with a FlashWeave correlation strength above 0.5 were kept as putative plasmid-host relationships.

### QUANTIFICATION AND STATISTICAL ANALYSIS

#### MAG polishing evaluation

As mentioned above, the final MAG collection was generated by binning the polished contigs, which were generated by metagenome assembly with MetaFlye, followed by three consecutive polishing steps (Medaka - long-read polishing, Polypolish - first round short-read polishing, and Polca - second round short-read polishing). To evaluate the effect of each polishing step, we generated “artificial” genomic bins containing the unpolished versions of the contig of the final MAG catalog for each one of the previous polishing steps (Flye, Medaka, Polypolish).

We evaluated polishing effects by looking at completeness and contamination values obtained by CheckM2. To assess statistical differences in the MAGs completeness and contamination values, we used SciPy library implementation of the Friedman test, followed by the Wilcoxon test on paired samples. Wilcoxon test on paired samples assessed significant differences between polishing steps (Flye - unpolished, Medaka - long-read polished, Polypolish - long-read + short-read polished, and Polca - long-read + 2x short-read polished), we evaluated if completeness between the polishing steps was higher (alternative=’greater’) and if contamination was lower (alternative=’less’). The final results were considered significant if the q-score obtained after Benjamini-Hochberg FDR correction was <0.05.

#### Taxonomic profiling and diversity analysis

To perform read-level analysis, we used short-read sequencing data from canid studies in public databases (up to February 2023) together with our short-read data. We kept the SingleM OTU tables generated by the 13 universal marker genes that target Bacteria and Archaea. For each OTU table, we filtered out OTUs that did not have a total count across all samples of at least 10 and that were not found in at least 2% of the samples. Moreover, we filtered out very low-abundant OTUs by removing any OTU with a mean relative abundance lower than 0.005%. Finally, we discarded samples that did not have at least 200 OTUs detected.

For each OTU table, we computed alpha diversity using the Shannon index and beta diversity using Bray-Curtis dissimilarity metrics (on log-transformed data). We obtained the final alpha and beta diversity objects by computing the median values for the selected marker genes. To assess statistical significance, we used Kruskal-Wallis for alpha diversity and PERMANOVA for beta diversity. We evaluated the effect of living environment, age, size, and sex. For each variable, if a specific category was only represented by a single study, the associated samples were filtered out to avoid batch effects in the statistical analysis.

#### Differential abundance analysis

Using SingleM species-level taxonomic profiles, we conducted differential abundance analysis using MaAsLin2^95^. For each metadata variable analyzed (living environment, age, size, and sex), *unknown* samples were discarded, and we adjusted for the other variables as relevant covariates. Features with a q-value < 0.1 were considered significant. Additionally, a Kruskal-Wallis test with Benjamini-Hochberg FDR correction (q < 0.05) was also conducted. Only taxa and functional features identified as significant by both methods and thresholds were considered robust associations and were further analyzed.

#### Comparison of canine long-read MAGs to representative genome assemblies

Using high-quality species-level genome assemblies, we compared our species representatives’ long-read MAGs to those from the public databases (GenBank or RefSeq). Firstly, we downloaded the reference genome for each species and ran CheckM2 on them to include only those species with >90% completeness and <5% contamination. We characterized them using the previously described workflow for counting tRNAs and identifying ribosomal genes.

To annotate the Mobilome function (COG category “X”) for both reference and long-read canine MAGs, we used eggNOG COG IDs and manually linked them to the COG X category according to NCBI (https://www.ncbi.nlm.nih.gov/research/cog/cogcategory/X/). Wilcoxon test on paired samples assessed if the count of mobilome gene functions was higher (alternative = ’greater’) in long-read species-level MAG here compared with representative genomes in public datasets. The final results were considered significant if the q-score obtained after Benjamini-Hochberg FDR correction was <0.05.

#### Extrachromosomal elements in canids

We mapped the short reads from a total of 406 dog samples from our Shanghai cohort, Berlin cohort, and selected external canid cohorts^7,9–11^ to the species-level MAGs and non-redundant ECEs. Reads were first quality filtered (trimmed to min. base quality of 25 using the ‘substrim’ strategy, and min. post-trim read size of 45) and then mapped to the dog reference genome with bwa-mem^96^ through NGLess^70^, keeping only hits with at least 45bp coverage at 90% identity. Reads mapping to the dog genome were then discarded and the remaining reads mapped to our catalog using the same software and filtering parameters. Using CoverM^85^, we obtained the “covered fraction” table and considered an EC element present if it was covered at least 80%.

Using the Chi-square test on the presence/absence table, we assessed differential prevalence patterns of ECEs across selected canid cohorts, considering several metadata variables: study, living environment, age, size, sex, and body condition. If the Chi-square test was significant, we ran a post-hoc Fisher’s Exact test for pairwise comparisons. As a final step, we performed multiple testing correction to all the combined results using the Benjamini-Hochberg FDR test.

Pairwise comparisons that included an unknown or unclassified group were filtered out for further analysis. For each ECE, we considered only the most significant metadata category driving the difference (the lowest corrected p-value). If that was ‘Study’, the difference is considered technical rather than biological, and is not further discussed.

##### Key resources table

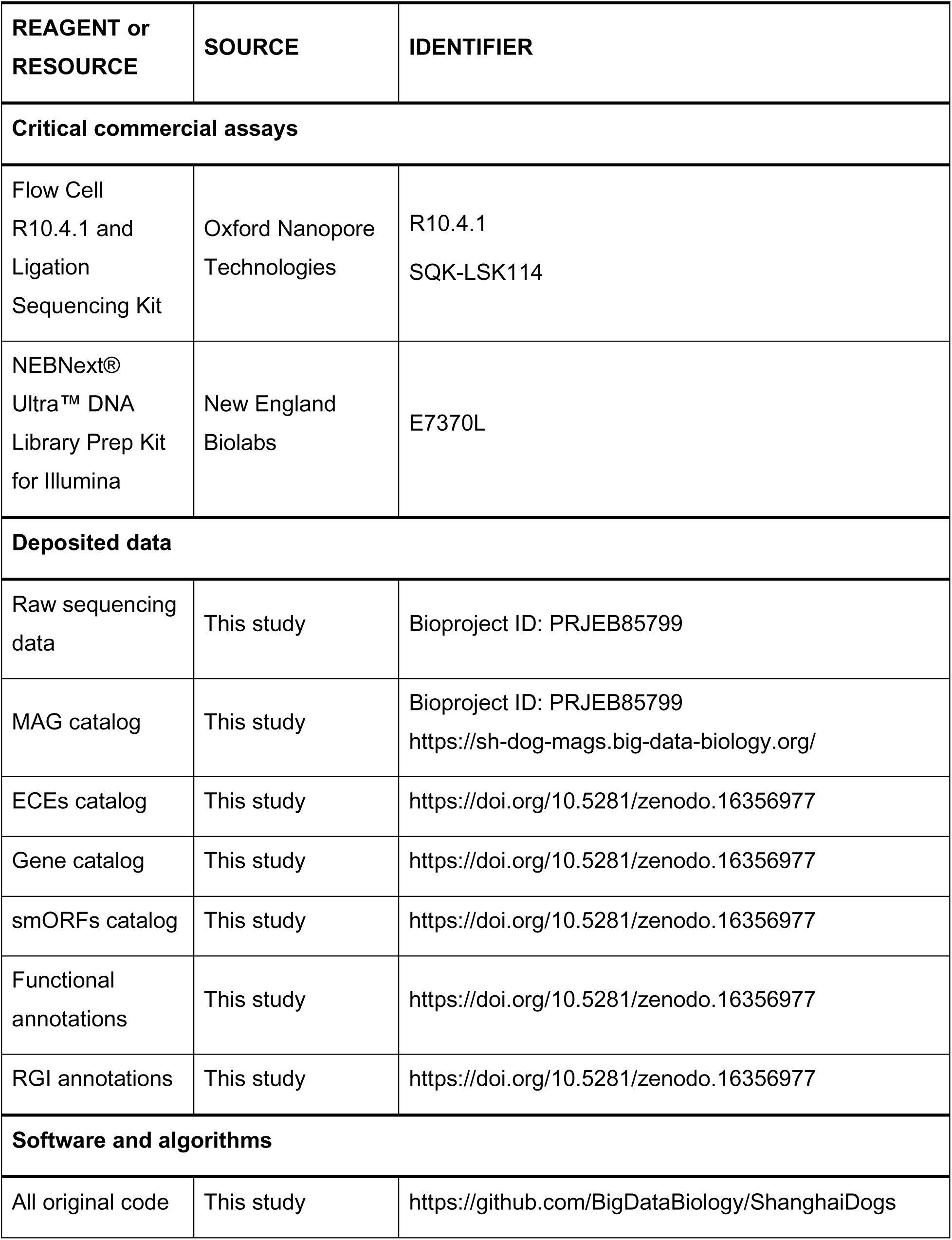
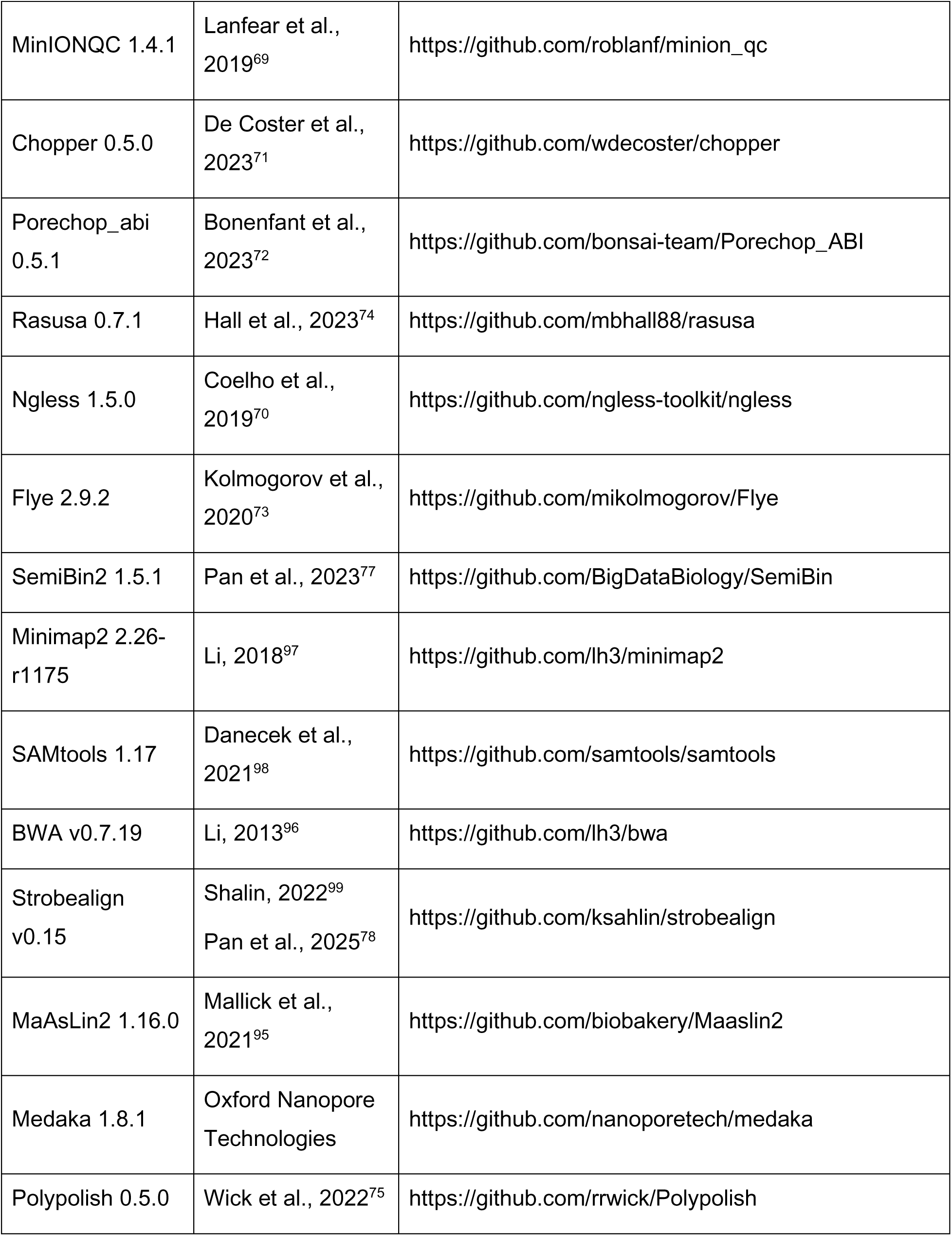
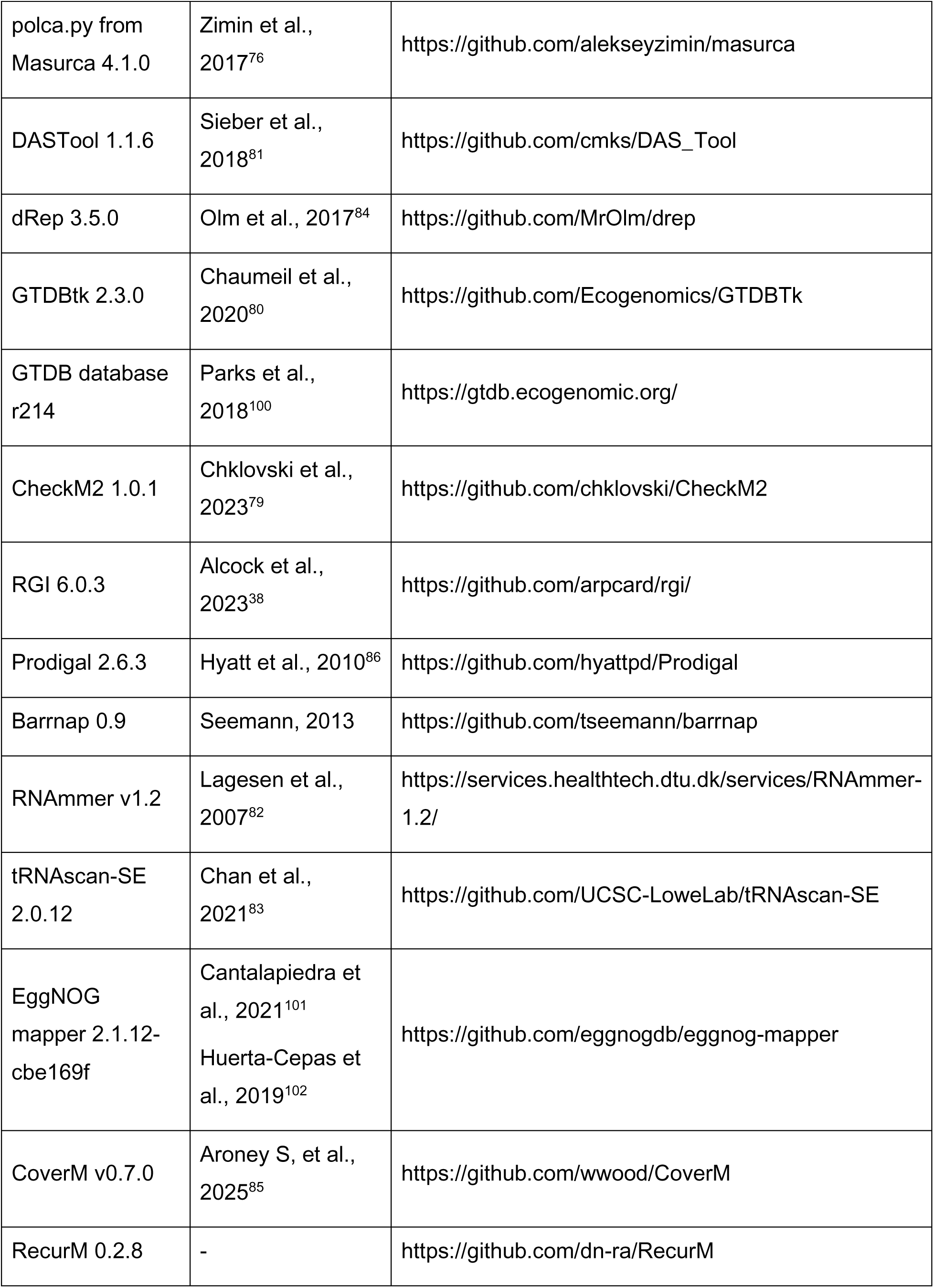
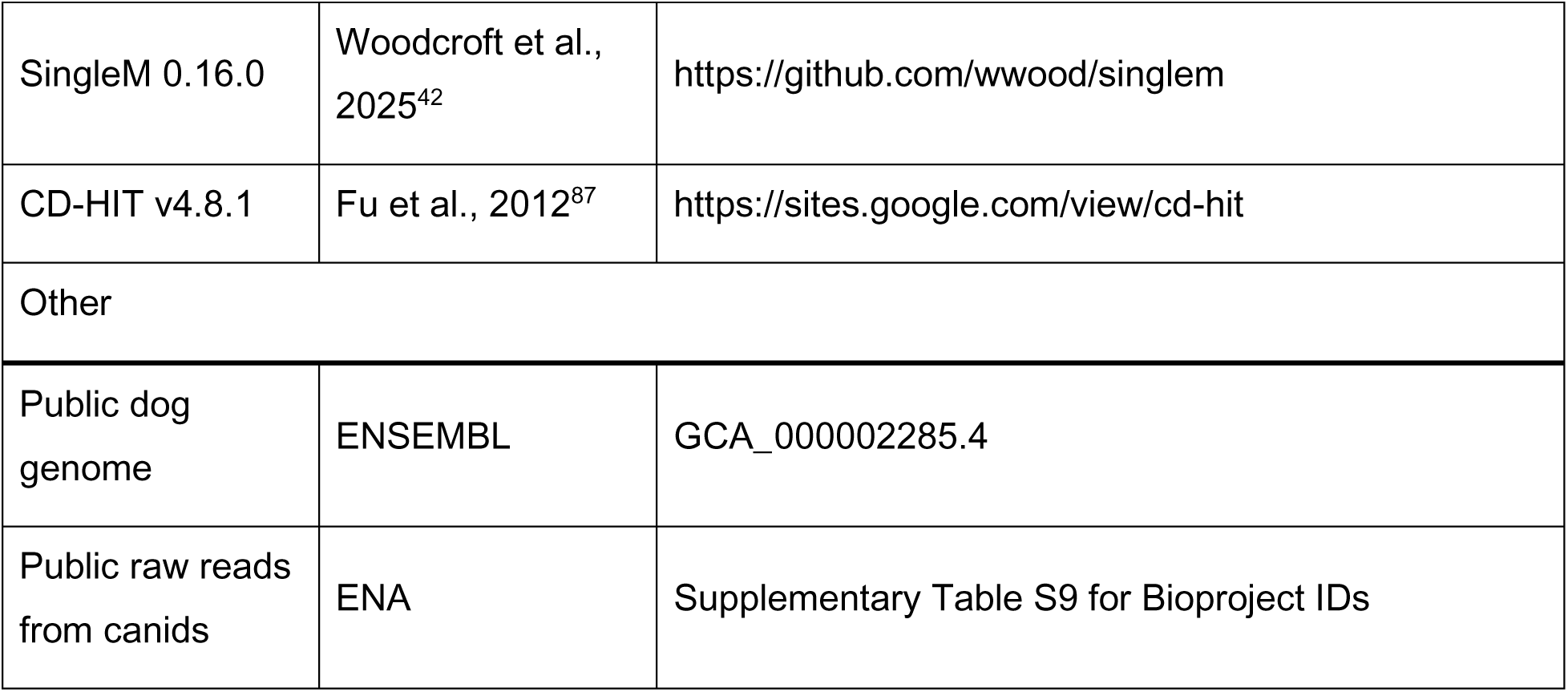

## Supporting information

Supplementary Table S1. Dog-associated information for the Shanghai cohort

Supplementary Table S2. Shanghai Dog owner questionnaire results

Supplementary Table S3. Shanghai Dog MAGs catalog metadata

Supplementary Table S4. Read mapping values of Canid datasets to Shanghai Dog catalogs

Supplementary Table S5. Comparison of representative species-level genome assemblies: Shanghai Dogs vs. reference

Supplementary Table S6. The 16S rRNA highest identity BLAST hit for species-level MAGs representing a novel species

Supplementary Table S7. Shanghai Dog circular extrachromosomal elements catalog metadata

Supplementary table S8. Metadata for the representative Canid samples: one sample per dog

Supplementary Table S9. Metadata for all the Canid datasets used

## Acknowledgements

The authors thank Alexandre Areias Castro (QUT) for assistance with sequence deposition, Chengkai Zhu (Fudan University) for support with logistics, Marion Draheim and Richard Lo-Man (Institut Pasteur Shanghai) for support with sample logistics, and the dog owners for their participation. Members of the Coelho group and the red herons group (QUT) are thanked for their feedback and suggestions throughout the project.

## Funding

National Natural Science Foundation of China (RFIS-I, 32250410281) (A.C.), the Australian Research Council (#FT230100724) (L.P.C.), the National Health and Medical Research Council of Australia (under the framework of JPI AMR, #2031902, SEARCHER) (L.P.C.), and the Deutsche Forschungsgemeinschaft (DFG, German Research Foundation) as part of the clinical research unit (CRU339): Food allergy and tolerance (FOOD@) – 409525714 (U.L.).

## Declaration of interests

A.Cu. is a partner at Nano1Health, SL, and has previously been invited by Oxford Nanopore Technologies (ONT) to participate in conferences. These activities did not influence the results or conclusions of this work. All other authors declare no competing interests.

## Author contributions

A.Cu. and L.P.C. designed the study. A.Cu. and F.G. collected the samples and the dog associated information. A.Cu., Y.D., A.Ch., S.P., X.-M.Z., and L.P.C. analyzed data. A.Cu., N.K., and L.P.C. visualized data. S.F., S.L., and U.L. provided the data from the Berlin dog cohort. A.Cu., X.-M.Z., and L.P.C. provided funding. L.P.C. supervised the project. A.Cu. wrote the first draft of the manuscript. All authors contributed to the revision of the manuscript prior to submission and approved the final version.

## SUPPLEMENTARY MATERIAL

**Supplementary Figure S1.**
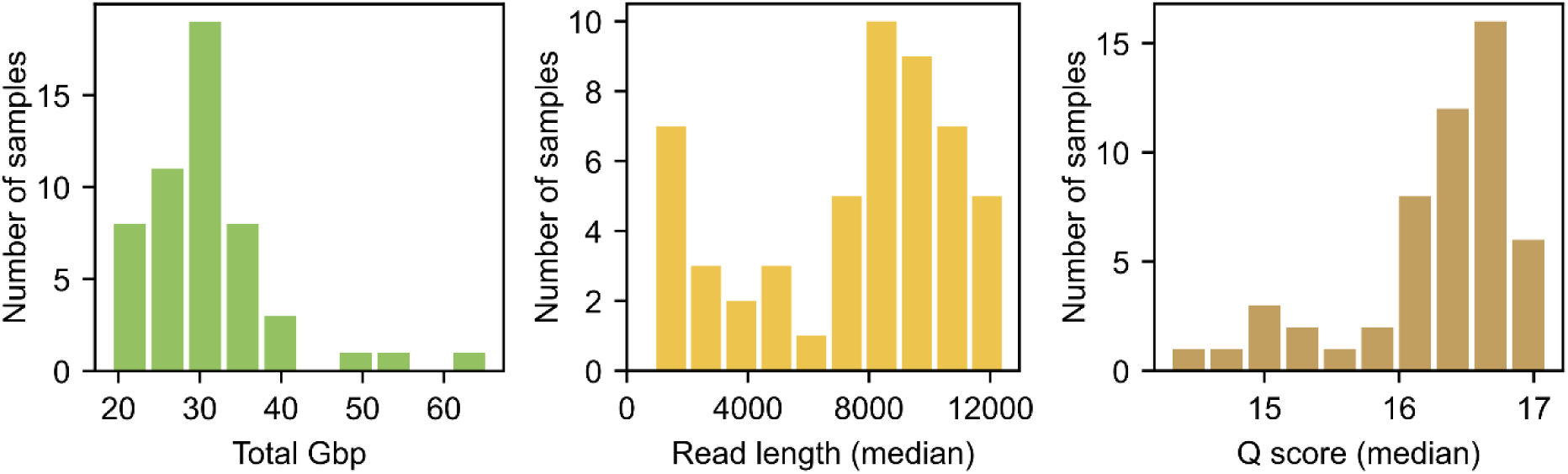
**Read-level summary statistics for ONT raw reads.**

**Supplementary Figure S2.**
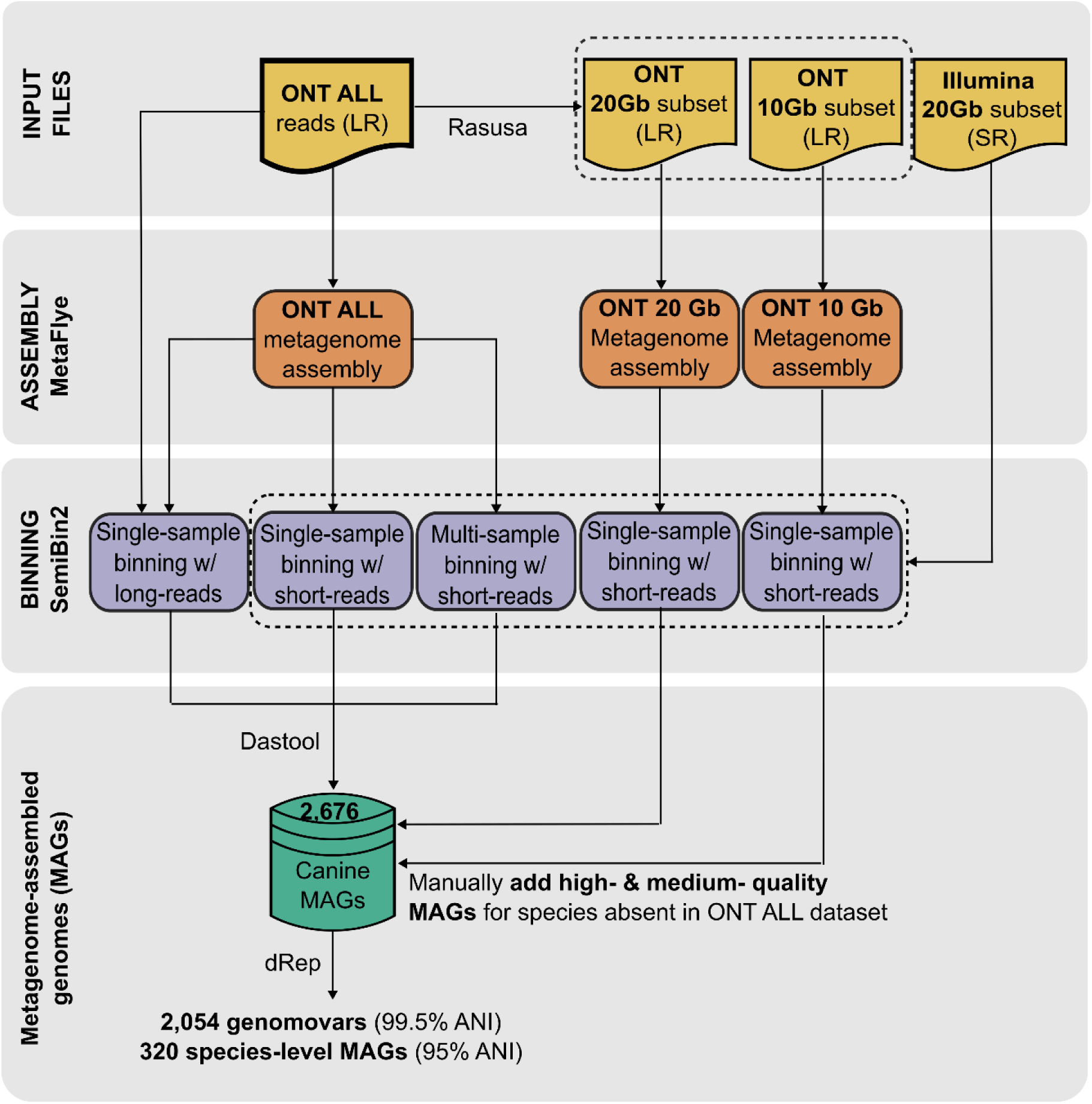
MAG generation pipeline detailed. Depending on the initial throughput of ONT reads, we created one or two subsets of data (20 Gbp and 10 Gbp). We performed metagenome assemblies on all the datasets (2-3 metagenome assemblies per sample). Using ‘ONT all’ assemblies, we performed three different binning strategies: single-sample binning using long reads for abundance estimation, single-sample binning using short reads for abundance estimation, and multi-sample binning using short reads for abundance estimation. Using subset assemblies, we performed single-sample binning with short reads for abundance estimation. We used Dastool to keep the best MAGs for each species from each sample. After assigning taxonomy using GTDB-tk, if subset assemblies contained a medium-or high-quality MAG representing a species absent in the MAG catalog, we manually added it.

**Supplementary Figure S3.**
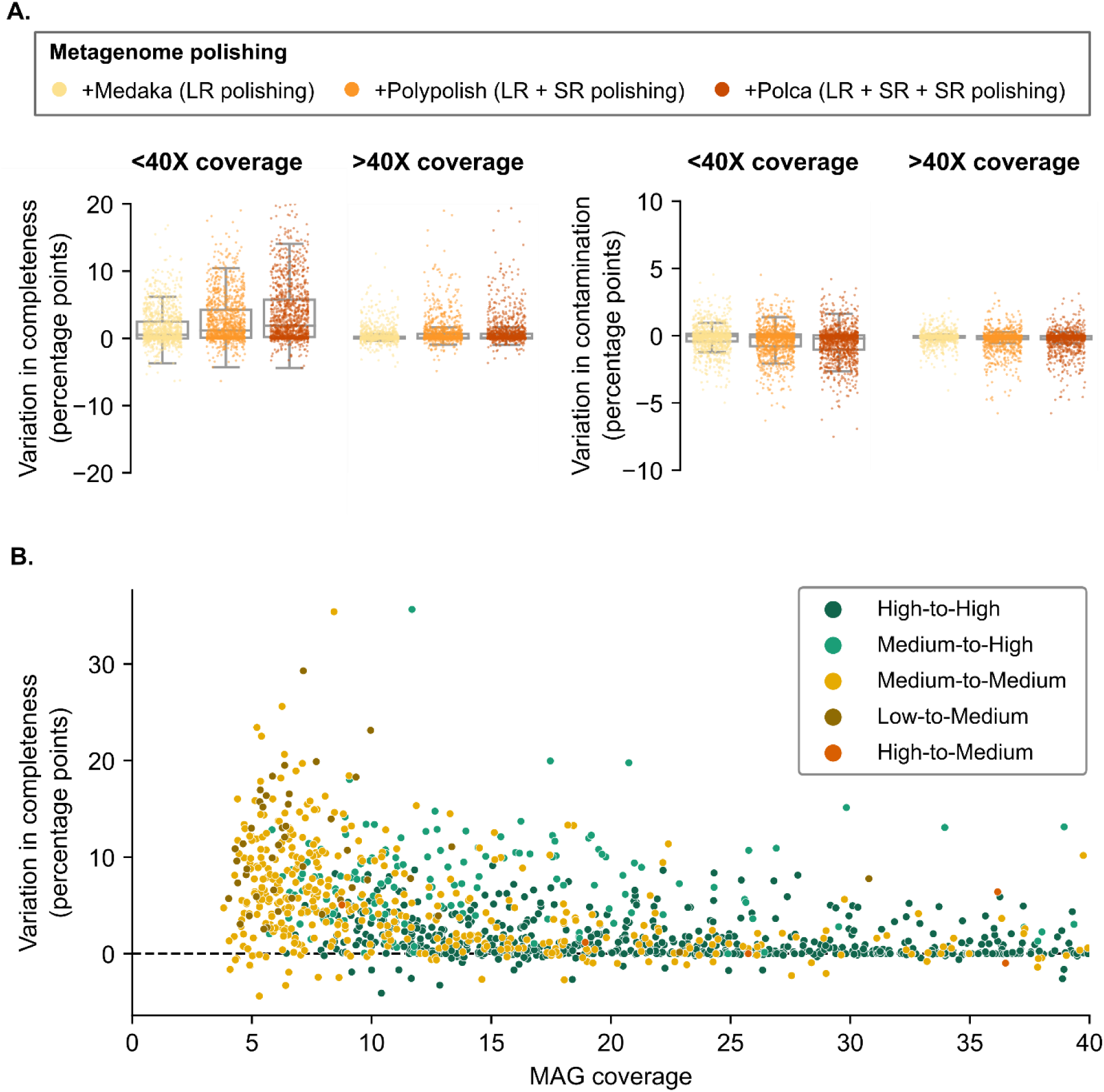
Polishing strategy evaluation in MAGs. **A)** Each polishing step, and especially those using short reads (SR), increased the quality of the MAGs by increasing completeness and reducing contamination (estimated with CheckM2). The effect was more significant for MAGs with lower coverage (<40X). **B)** Unpolished to (most) polished MAG evolution for low-covered MAGs: we can observe a general increase in completeness values, which may lead to transforming medium-quality MAGs to high-quality MAGs.

**Supplementary Figure S4.**
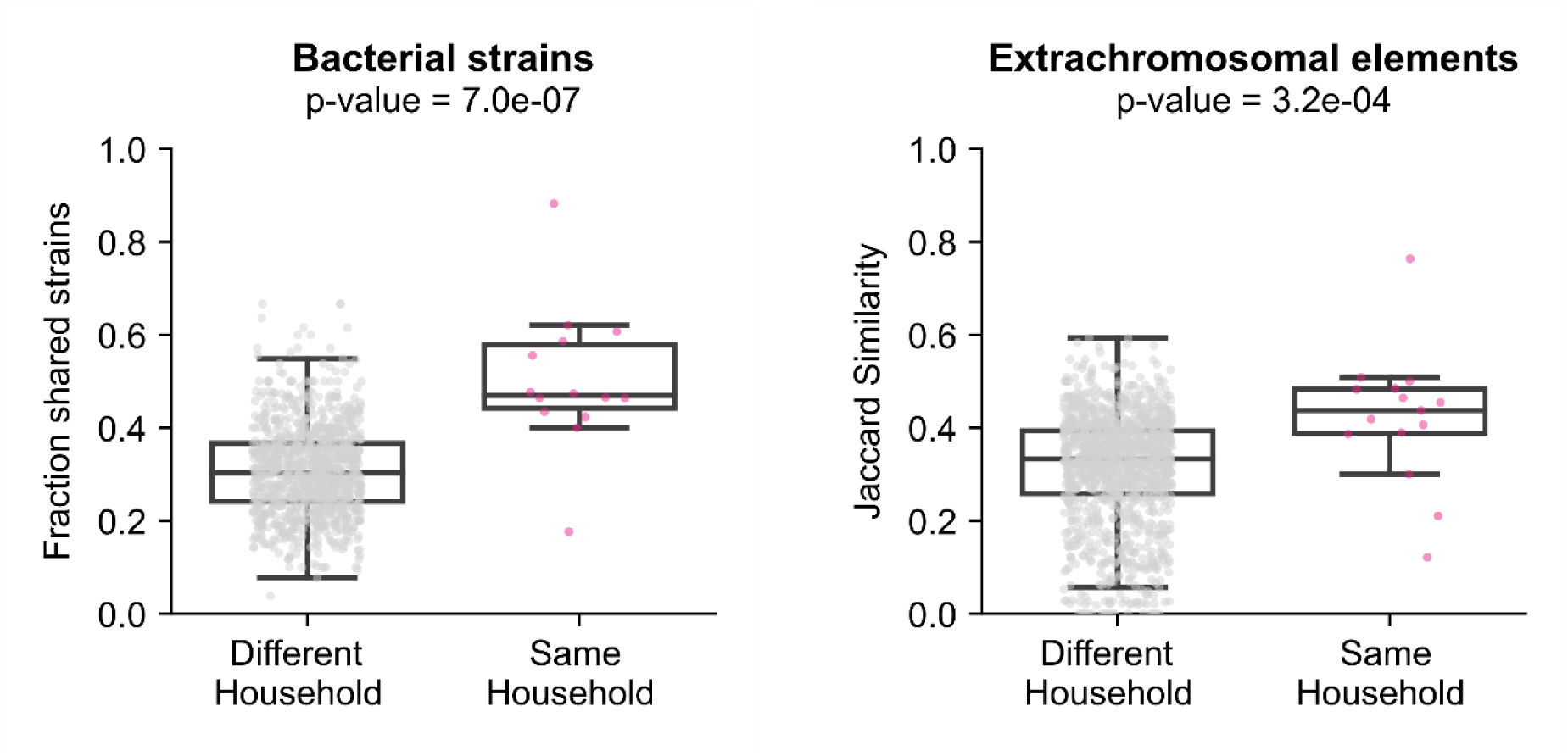
Strain and extrachromosomal elements sharing within the Shanghai Dog cohort. We performed pairwise comparisons (Mann–Whitney U test, one-sided test) between dogs within the same households vs. between households, and found significantly more shared strains (left) and EC elements (right) in dogs sharing households.

**Supplementary Figure S5.**
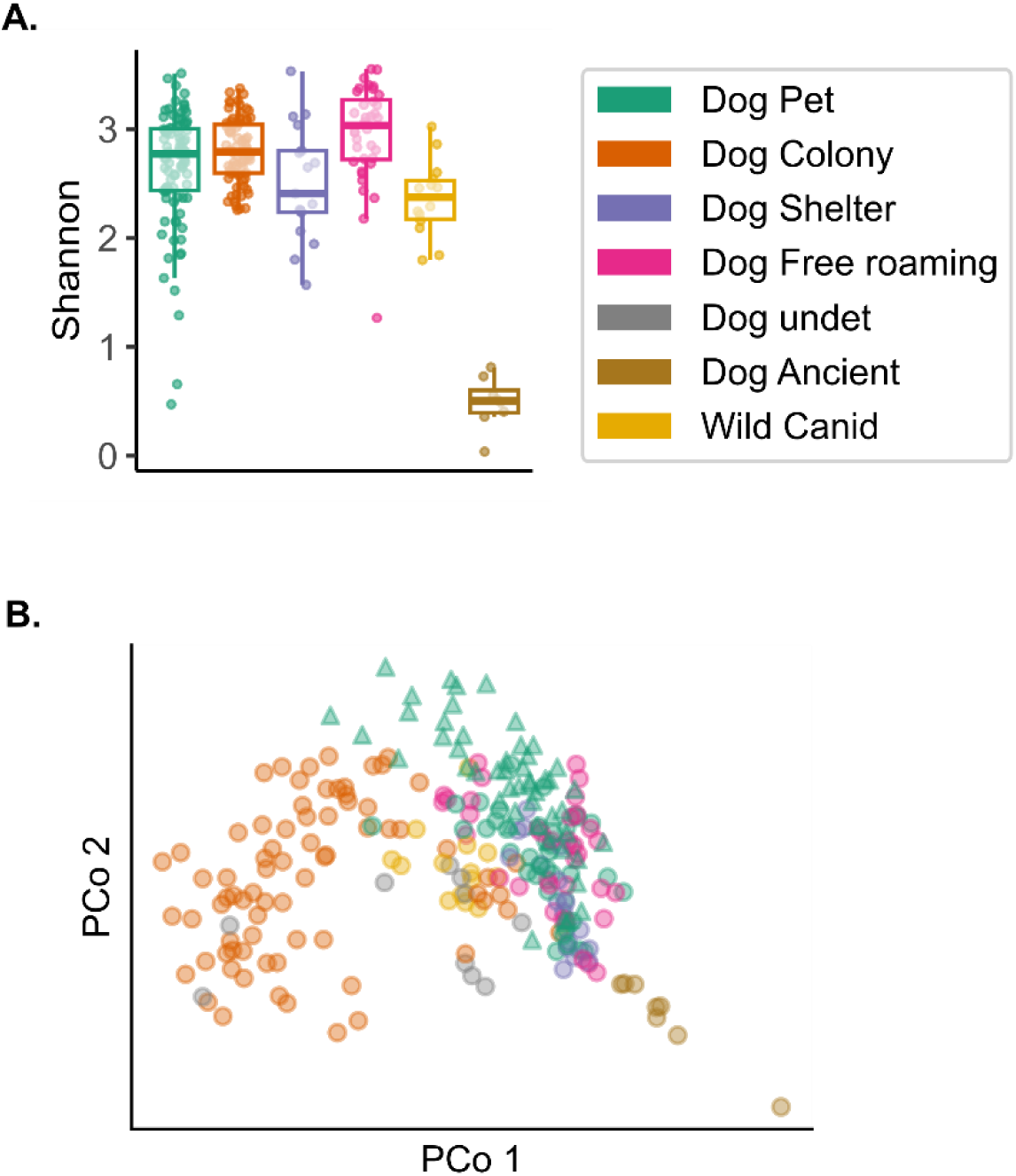
Living environment effect on the dog gut microbiome, including all Canid groups. Analogous to Fig. 5A-B, here, we also included free-roaming dogs, ancient canid samples, and unclassified canid samples. The colored legend applies to the whole figure. **A)** Boxplots representing alpha diversity (Shannon index). **B)** PCoA plot representing beta diversity (Bray-Curtis on log-transformed data). Green triangles indicate pet dogs in this study.

**Supplementary Figure S6.**
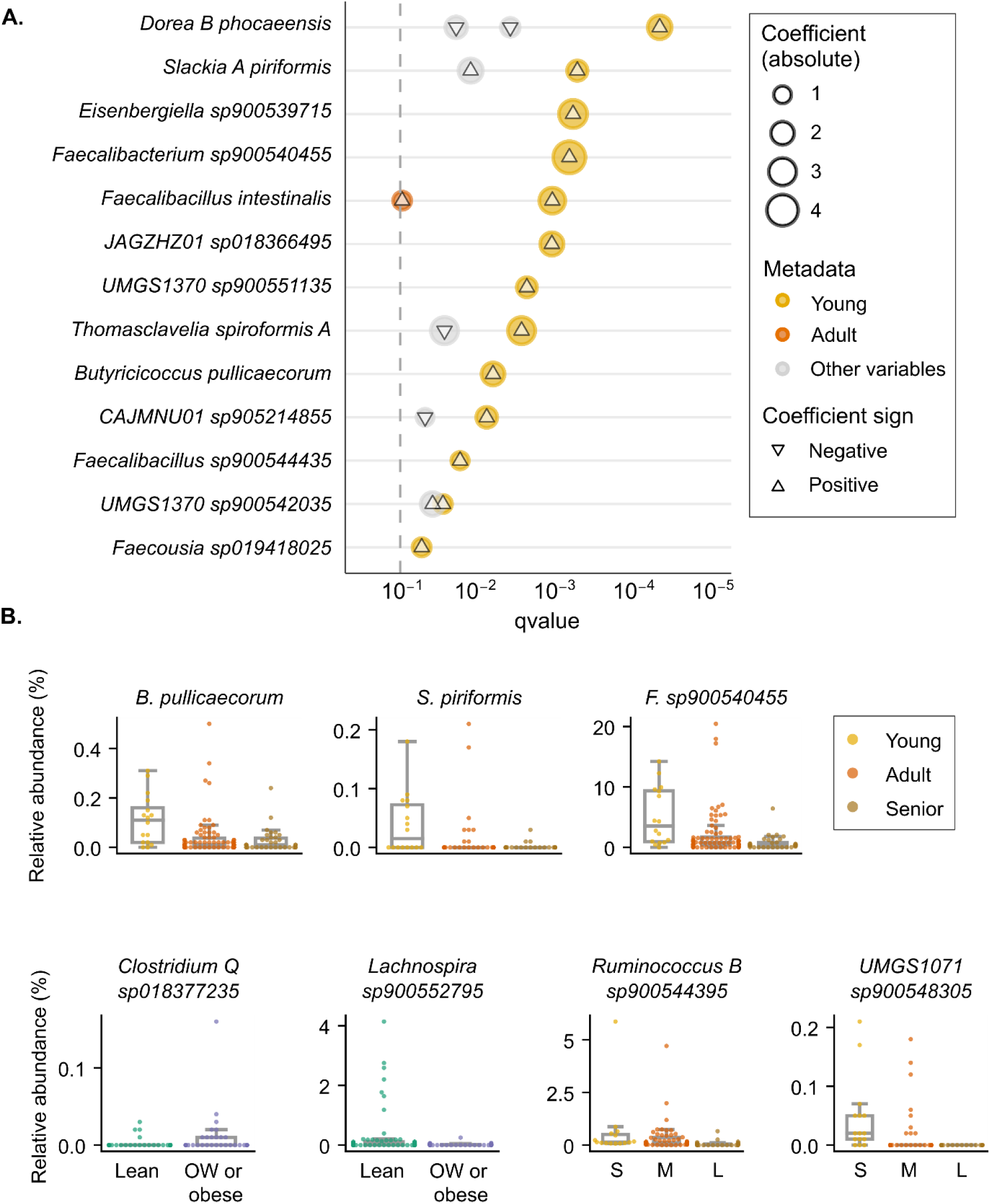
Differential abundance analysis results by age, size, and body condition. **A)** Differentially abundant bacterial species associated with dog age (concordant results, Maaslin2 and Kruskal-Wallis). In the y-axis, the significant taxa, and in the x-axis, their associated q-value in Maaslin2. We used senior dogs as the reference group in Maaslin2, so circles are colored according to the group that differs from them, and they are scaled in concordance with the absolute coefficient value. Grey circles represent significant differences found in that species when comparing groups in another variable, rather than age (age being the most significant variable). Triangles within the circles indicate if the coefficient is negative (inverted triangle, less abundant species) or positive (more abundant). **B)** Boxplots of total relative abundances (%) of selected differentially abundant bacterial species, considering age (young, adult, and senior); body condition (lean, overweight/obese); and size (small, medium, large).

## Supplementary tables

**Supplementary Table S1. Dog-associated information for the Shanghai cohort (n=51).** A summarized version of the most relevant questionnaire results, with appropriate metadata categories considering the questionnaire output (with the latest information regarding animal health status).

**Supplementary Table S2. Shanghai Dog owner questionnaire results (n=107).** From questionnaire to final sample collection, several days to weeks might have passed, so questionnaire results regarding health status might slightly differ from the final metadata (Supplementary Table S1).

**Supplementary Table S3. Shanghai Dog Metagenome-assembled genomes catalog metadata.** Includes taxonomic classification, quality information, and other descriptive characteristics of the MAGs.

**Supplementary Table S4. Read mapping values of Canid datasets to Shanghai Dog catalogs.** Values include mapping to the whole MAG catalog (‘aligned’), to the representative species MAGs (‘aligned_sp’), and to the catalog and extrachromosomal elements (‘aligned_EC’). It also includes read mapping summary statistics by study, and by living environment.

**Supplementary Table S5. Comparison of representative species-level genome assemblies: Shanghai Dogs *vs.* reference.** For each bacterial species, we compared the representative genome assembly from Shanghai Dog cohort to that found in a public database (RefSeq or GenBank). Only species that had a high-quality genome assembly (>90% completeness and <5% contamination) were included. Comparison included the counts of: contigs, ribosomal genes, and mobilome COG hits (COG category X), and statistical differences were assessed using Wilcoxon pairwise test.

**Supplementary Table S6. The 16S rRNA highest identity BLAST hit for species-level MAGs representing a novel species.** The table only includes high-quality MAGs that have at least one of their 16S rRNA hits with >99% identity to 16S rRNA NCBI databases.

**Supplementary Table S7. Shanghai Dog circular extrachromosomal elements catalog metadata (n=185).** The table includes basic descriptive information (size, sample origin, and category of the ECE). Uncategorized elements might include genetic markers for both viruses and plasmids, or may lack identifiable markers. Finally, for 52 of them, we computationally predicted the putative bacterial host.

**Supplementary table S8. Metadata for the representative Canid samples: one sample per dog.** This metadata includes Shanghai Dogs, and external shotgun metagenomics studies from: public datasets (up to February 2023); a new pet dog cohort in Germany. For longitudinal studies, we chose a single representative sample per dog based on: the baseline diet (*e.g.,* in dietary intervention studies); or the ‘healthiest’ status (*e.g.,* absence of clinical signs after treatment, in cases of chronic enteropathies).

**Supplementary Table S9. Metadata for all the Canid datasets used.** For external public datasets, it includes dog shotgun metagenomics studies up to February 2023. Finally, it also includes metadata for a pet dog cohort in Germany.

## Notes

https://sh-dog-mags.big-data-biology.org/

https://doi.org/10.5281/zenodo.16356977

### Competing Interest Statement

Anna Cusco is a partner at Nano1Health, SL, and has previously been invited by Oxford Nanopore Technologies (ONT) to participate in conferences. These activities did not influence the results or conclusions of this work. All other authors declare no competing interests.

